# *Trpv1*-dependent *Cacna1b* gene inactivation reveals cell-specific functions of Ca_V_2.2 channels *in vivo*

**DOI:** 10.1101/2025.05.09.652131

**Authors:** Remy Y. Meir, Martin S. Sisti, Arturo Andrade, Diane Lipscombe

## Abstract

Voltage-gated Ca_V_2.2 channels underlie the N-type current, and they regulate calcium entry at many presynaptic nerve endings to control transmitter release. A role for Ca_V_2.2 channels has been well-established in the transmission of pain information using pharmacological and global gene inactivated mouse models. However, investigation of the cell-specific actions of Ca_V_2.2 channels would benefit from the availability of cell-restricted knockout mouse models and particularly in dissecting behavioral responses that depend on Ca_V_2.2 channel activity. Here, we show the importance of Ca_V_2.2 channels in *Trpv1*-lineage neurons in behavioral responses to sensory stimuli using Cre-dependent inactivation of the *Cacna1b* gene. Our work shows the cell-type specificity of Ca_V_2.2 channels in mediating rapidly developing heat hypersensitivity and the utility of Cre-dependent inactivation of *Cacna1b* to discern cell-specific Ca_V_2.2 channel functions.

## INTRODUCTION

Voltage-gated Ca_V_2.2 channels regulate calcium entry at many presynaptic nerve endings to control transmitter release. At some synapses, Ca_V_2.2 channels co-regulate synaptic transmission with Ca_V_2.1 channels whereas, at other synapses including in the brain, and in sympathetic and sensory neurons, Ca_V_2.2 channels are the dominant source of calcium that triggers transmitter release^1,2^. Ca_V_2.2 channels are well known targets of numerous neuromodulators and drugs that down regulate synaptic efficacy via G protein coupled receptor activation^3^. At sensory nerve endings in skin, Ca_V_2.2 channels play a key role in neuroimmune signaling during neuroinflammation^4,5^.

The availability of highly specific toxins for Ca_V_2.2, ω-CgTx-GVIA and ω-CgTx-MVIIA, have helped define functions of Ca_V_2.2 channels in certain cells and tissues^6–9^. *Cacna1b* global knockout mouse models have linked Ca_V_2.2 channel activity to several behaviors, including the threshold for withdrawal responses to sensory stimuli, overall activity, Rapid Eye Movement (REM) sleep and vigilance^4,5,10–12^. But conditional *Cacna1b^-/-^* mouse strains would benefit studies aimed at isolating cell-specific actions of Ca_V_2.2 channels and dissecting their contributions to specific neuronal circuits and behaviors. In addition, such models would improve analyses aimed at discerning the contribution of Ca_V_2.2 channels to specific behaviors as compared to global *Cacna1b^-/-^* mouse models which all exhibit a strong hyperactivity phenotype^10,11,13,14^.

Here we use a *Cacna1b^fl/fl^* floxed mouse strain for Cre-dependent inactivation of *Cacna1b* in *Trpv1*-lineage neurons. We show the dual function of Ca_V_2.2 channels in *Trpv1*-expressing, noxious heat-sensing neurons; controlling the efficacy of transmission from peripheral to central sites and triggering heat hypersensitivity in skin associated with neuroinflammation. We also show that Ca_V_2.2 channels contribute to paw withdrawal responses to cold, as well as heat stimuli consistent with their importance in triggering transmitter release at presynaptic sites.

## RESULTS

### Ca_V_2.2 protein levels reduced in DRG but not in brain of conditional knockout mice

*Cacna1b^fl/fl^* (WT-fl) and *Trpv1^Cre/Cre^* (WT-cre) breeding pairs were used to generate *Cacna1b^fl/fl^*/*Trpv1^Cre^*^/-^ (cKO) mice lacking functional Ca_V_2.2 channels in *Trpv1*-lineage neurons (Fig. 1A). cKO mice were viable and fecund and, by these metrics, similar to WT-fl littermate controls (Table 1). *Trpv1* is expressed primarily in sensory neurons of dorsal root ganglia (DRG) including in heat and cold sensing subtypes^15–17^. We used polyclonal anti-Ca_V_2.2 antibodies^10,18,19^, to compare levels of Ca_V_2.2-specific signals in protein preparations from brain and DRG of cKO, *Cacna1b^-/-^* (global knockout; gKO) and WT-fl mice (Fig. 1B). Polyclonal anti-Ca_V_2.2 target an epitope located on the intracellular loop between domains II and III of Ca_V_2.2. These polyclonal antibodies recognizes a number of protein bands covering a range of molecular weights in protein preparations of the brain and DRG (Fig. 1B) consistent with previously published findings from our group using polyclonal anti-Ca_V_2.2 in native tissue^19,20^. The use of gKO tissue was therefore essential to verify the identity of Ca_V_2.2-specific bands in brain (a single band at ∼280 kDa) and in DRG (two bands at ∼280 kDa and ∼230 kDa). In cKO and WT-fl control brain protein preparations, the intensity of Ca_V_2.2-specific bands were similar (Fig. 1B), as expected given the very low level of *Trpv1* expression in mouse brain^17,21,22^. By contrast, in DRG which contains relatively high abundance of *Trpv1*-lineage DRG neurons^23,24^, the intensity of Ca_V_2.2-specific bands from cKO were ∼50% of WT-fl controls (Fig. 1B). These data are as expected for selective inactivation of *Cacna1b* in *Trpv1*-lineage neurons and they highlight the importance of using gKO tissue to identify Ca_V_2.2-specific signals using polyclonal antibodies in native tissue^25^.

**Figure 1.**
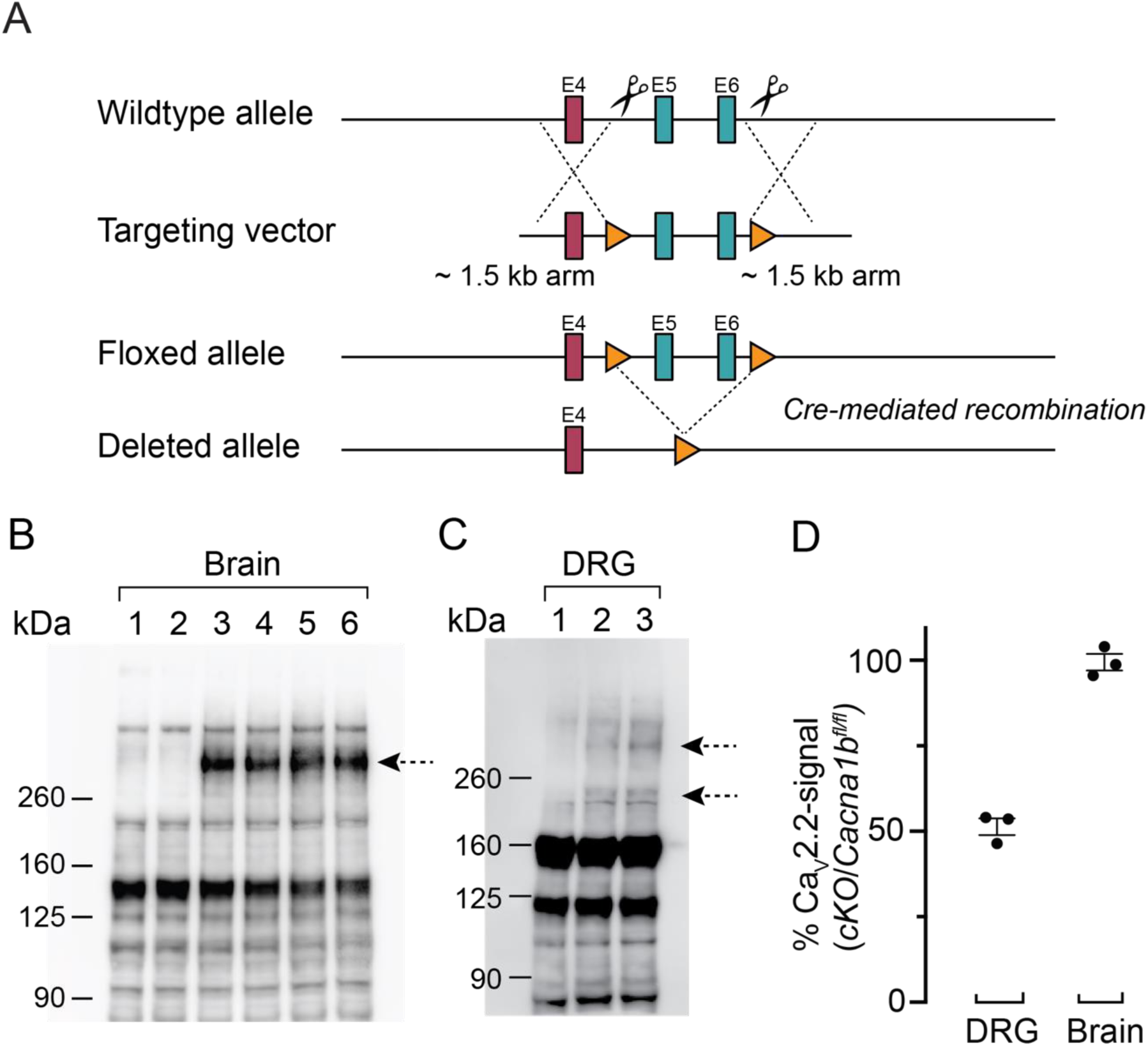
*Cacna1b* gene targeting strategy and Ca_V_2.2 protein analysis of brain and dorsal root ganglia. Gene targeting strategy (***A***). CRISPR-Cas9 insertion of loxP sites in introns flanking exons 5 and 6 of *Cacna1b* (mm10 chr2:24,603,887-24,763,152). ***B, C*,** Western analyses of protein preparations from brain (***B***: lanes 1-6) and DRG (***C***: lanes 1-3) of *Cacna1b*^-/-^ (gKO; ***B***, lanes 1, 2; ***C***, lane 1), *Cacna1b*^fl/fl^/*Trpv1^Cre/-^* (cKO; ***B***, lanes 3, 4; C, lane 2) and *Cacna1b*^fl/fl^ (WT-fl, ***B***, lanes 5, 6; ***C***, lane 3) mice. Anti-Ca_V_2.2 polyclonal antibody (ACC-002, Alomone Labs) identified one prominent Ca_V_2.2-specific band in brain at ∼280 kDa in cKO and WT-fl; and two Ca_V_2.2-specific bands at ∼280 and ∼240 kDa in cKO and WT-fl DRG. ***D***, Intensity of Ca_V_2.2- specific signals in brain and DRG of cKO relative to WT-fl are plotted for 3 experiments. The average band intensity (in arbitrary units) for brain was WT-fl 95.3 ± 5.7 and cKO 101.7 ± 2.3 and were not significantly different from control (WT-fl | cKO: *t* = 0.9759 *p* = 0.3668; Student’s t-test).

**Table 1.**
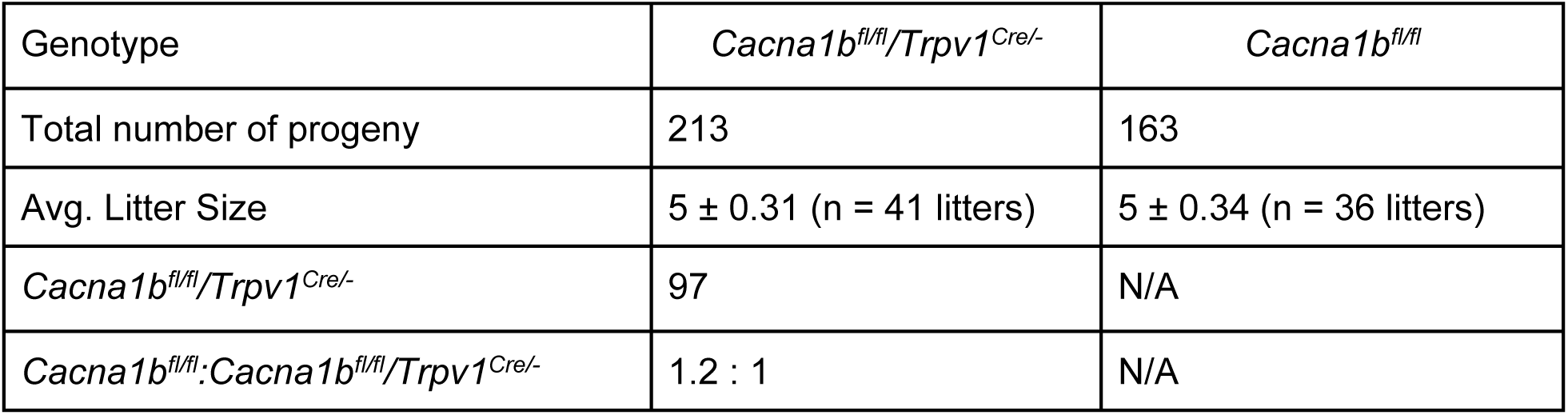
Cacna1b^fl/fl^*/Trpv1^Cre/-^* mice are viable and fecund and show no differences respect to WT-fl littermate controls.

### Whole cell Ca_V_2.2 currents absent in *Trpv1*-positive DRG neurons of cKO mice

To confirm selective loss of Ca_V_2.2 protein in *Trpv1*-lineage neurons of DRG, we compared whole cell Ca_V_ currents in DRG neurons isolated from cKO, WT-cre and WT-fl mice. Recordings were intentionally biased toward small diameter neurons which contain the highest percentage of *Trpv1*-expressing neurons^18^ (Fig. 2A). The average size of neurons in this analysis, based on measurements of total cell capacitance, was not statistically different among the three genotypes (Fig. 2A: WT-fl = 15.3 ± 0.88 pF, cKO = 16.8 ± 1.10 pF and WT-cre = 15.2 ± 0.65 pF; WT-fl | cKO | WT-cre: *H* = 0.105, *p* = 0.9489; Kruskal-Wallis). Ca_V_ currents were recorded from a total of 107 DRG neurons in response to step depolarizations applied in 5 mV increments, from a holding potential of -80 mV (2 mM Ca^2+^ was used as the charge carrier; Fig. 2B). Total Ca_V_ current densities recorded from WT-cre and WT-fl control DRG neurons were not statistically distinguishable (Fig. 2C; WT-cre | WT-fl: *t* = -0.446, *p* = 0.6574; Student’s t-test) so we used WT-cre controls for all electrophysiology experiments due to ability to label *Trpv1*-lineage neurons.

**Figure 2.**
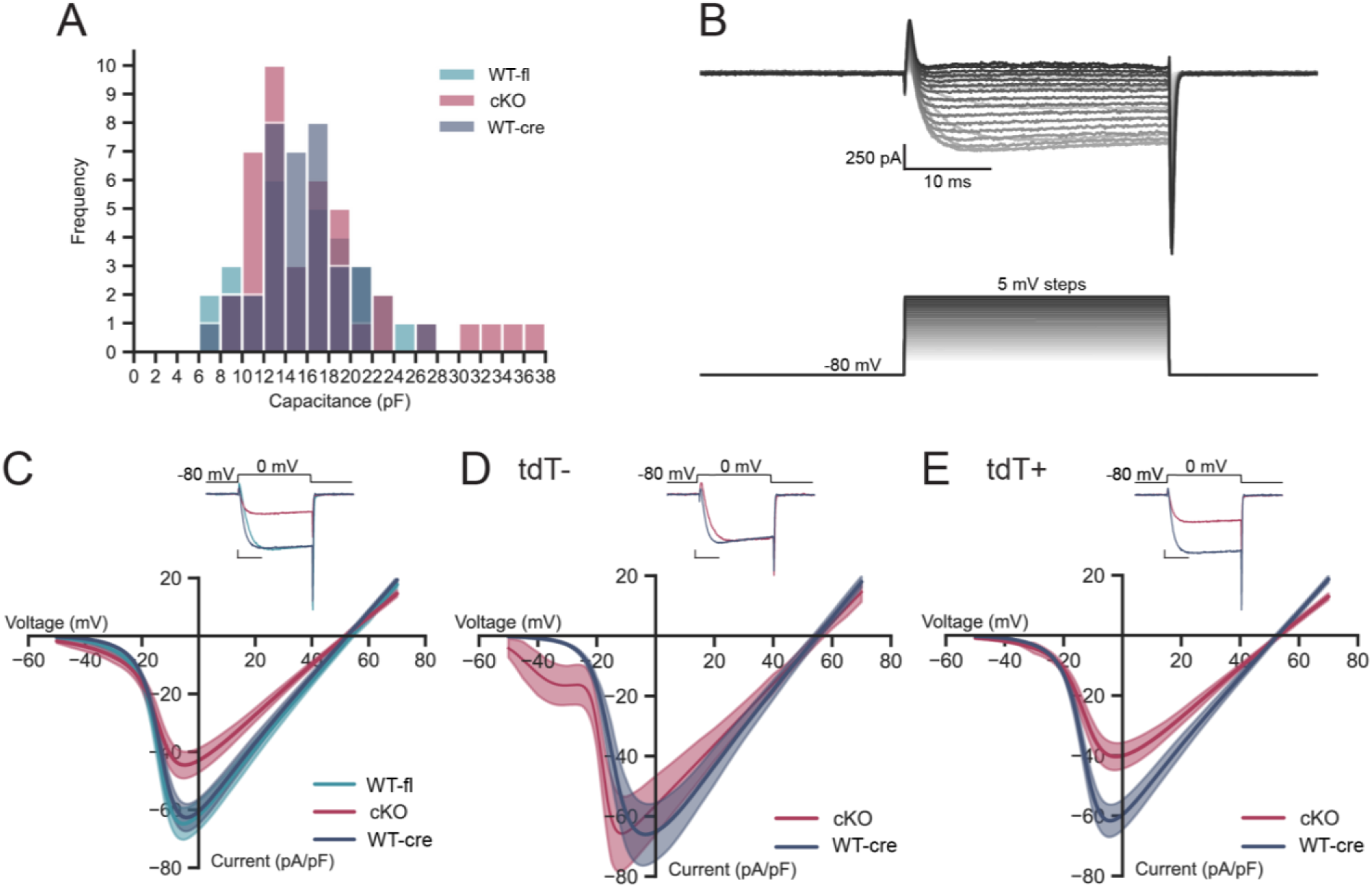
Whole cell voltage-gated Ca_V_ currents are reduced in *Trpv1*-positive neurons from *Cacna1b^fl/fl^/Trpv1^Cre/-^* cKO mice. Ca_V_ current-voltage relationships recorded from acutely dissociated DRG neurons from *Cacna1b^f^*^l/fl^ (WT-fl, light blue; *n* = 31), *Trpv1^Cre/Cre^* (WT-cre, dark blue; *n* = 35) and *Cacna1b^fl/fl^/Trpv1^Cre/-^* (cKO, red; *n* = 41). ***A***, Whole cell capacitance measurements from DRG neurons from WT-fl, (light blue; 15.3 ± 0.88 pF), WT-cre (dark blue; 15.2 ± 0.65 pF) and cKO (red; 16.8 ± 1.10 pF) mice were not statistically distinguishable (*H* = 0.105, *p* = 0.9489; Kruskal-Wallis). ***B***, A series of Ca_V_ currents from a single neuron elicited by depolarizing steps from a holding potential of -80 mV to voltages between -50 mV and + 70 mV in 5 mV increments. ***C****, **D**, **E***, Average Ca_V_ current densities (pA/pF) plotted as a function of the test voltage for all neurons from each of the three genotypes (***C***), *Trpv1*-positive (tdT+; ***D***), and *Trpv1*-negative (tdT−; ***E***) neurons. Insets for each panel show representative Ca_V_ current recordings evoked by steps to 0 mV from -80 mV; scale bars correspond to 200 pA and 10 ms (**C, D, E**). ***C***, Total inward Ca_V_ current of current-voltage relationships generated from recordings from DRG neurons of WT-fl (light blue) and WT-cre (dark blue) mice are not statistically different (*t* = -0.446, *p* = 0.6574; Student’s t-test). Viral delivery of tdTomato in cKO and WT-cre is used to identify *Trpv1*-lineage neurons (tdT+) (***D***, ***E***). ***D***, In tdT− neurons, the total Ca_V_ current calculated as the integral of current-voltage relationships is not statistically different between cKO (red, *n* = 9) and WT-cre (blue, *n* = 8) (WT-cre | cKO: *t* = -0.575, *p* = 0.5735; Student’s t-test). ***E***, In tdT+ neurons, the total Ca_V_ current from current-voltage relationships is statistically different between cKO (red, *n* = 32) and WT-Cre (blue, *n* = 27) recordings (WT-cre | cKO: *t* = 2.911, *p* = 0.005; Student’s t-test). In cKO/ tdT− neurons a prominent low threshold Ca_V_3-type current is evident. Ca_V_3 currents are a prominent feature of non-*Trpv1*-expressing mechanosensory DRG neurons which have slightly larger cell diameters compared to *Trpv1*-lineage neurons^31^ (as seen in the capacitance plot (***A***) and see^18^). Ca_V_ current-voltage relationships show average (solid line) ± SEM (shaded area) values together with the average fitted curves using one (***C, E***) or the sum of two (***D***) Boltzmann-linear line functions. Average parameters calculated from the fitted curves of individual cells.

We used adeno-associated viral (AAV) delivery of Cre-dependent tdTomato (tdT) to label *Trpv1*- lineage neurons in WT-cre and cKO mice and compared Ca_V_ currents in non-*Trpv1*-lineage (non-Cre expressing; tdT−) and *Trpv1*-lineage (Cre-expressing; tdT+) neurons (Figs. 2D, 2E). In cKO mice, total Ca_V_ current densities in tdT− neurons (*n* = 9) were statistically indistinguishable from tdT− neurons of WT-cre controls (*n* = 8) (Fig. 2D; *t* = -0.571, *p* = 0.5735; Student’s t-test). By contrast, in cKO mice, total Ca_V_ current densities in tdT+ neurons (*n* = 32) were significantly lower compared to tdT+ neurons of WT-cre controls (*n* = 27) (Fig. 2E; *t* = 2.911, *p* = 0.0051; Student’s t-test). These data show there is selective loss of Ca_V_ currents in tdT+, but not tdT− neurons, of cKO mice and they confirm selective inactivation of *Cacna1b* in *Trpv1*-lineage neurons. There is a low threshold Ca_V_3 current in recordings from tdT−, but not in tdT+ neurons of cKO mice (Fig. 2D). This current corresponds to *Cacna1h*-expressing mechanoreceptors which do not express *Trpv1*^23,24,26–29^ and which are slightly larger than *Trpv1*-expressing neurons^18^ (see Fig. 2A).

ω-Conotoxin GVIA (ω-CgTx-GVIA) is a potent, selective inhibitor of Ca_V_2.2 channels. We compared the effects of ω-CgTx-GVIA on Ca_V_ currents in tdT− and tdT+ neurons of WT-cre and cKO DRG. ω-CgTx-GVIA inhibited Ca_V_ currents in tdT− neurons from WT-cre (∼50%) and cKO (∼42%) mice (Fig. 3A, tdT−/WT-cre, control | ω-CgTx: *n* = 6, *t* = -2.573, *p* = 0.0277; Fig. 3B, cKO, control | ω-CgTx: *n* = 5, *t* = -2.761, *p* = 0.0246; Student’s t-test). ω-CgTx-GVIA also reduced Ca_V_ currents in tdT+/WT-cre neurons (∼50%) (Fig. 3C, tdT+/WT-cre, control | ω-CgTx: *n* = 6, *t* = - 3.584, p = 0.0050; Student’s t-test), whereas, Ca_V_ current densities in tdT+/cKO neurons were only slightly smaller in the presence of ω-CgTx-GVIA (Fig. 3D; tdT+/cKO, control | ω-CgTx: *n* = 6, *t* = -2.101, *p* = 0.082; Student’s t-test). The slight reduction in maximum Ca_V_ current densities in cKO neurons in the presence of ω-CgTx-GVIA, as compared to WT-cre controls, likely represents Ca_V_1 current run-down during whole cell recording (Fig. 3C; also see supplementary data)^30,31^.

**Figure 3.**
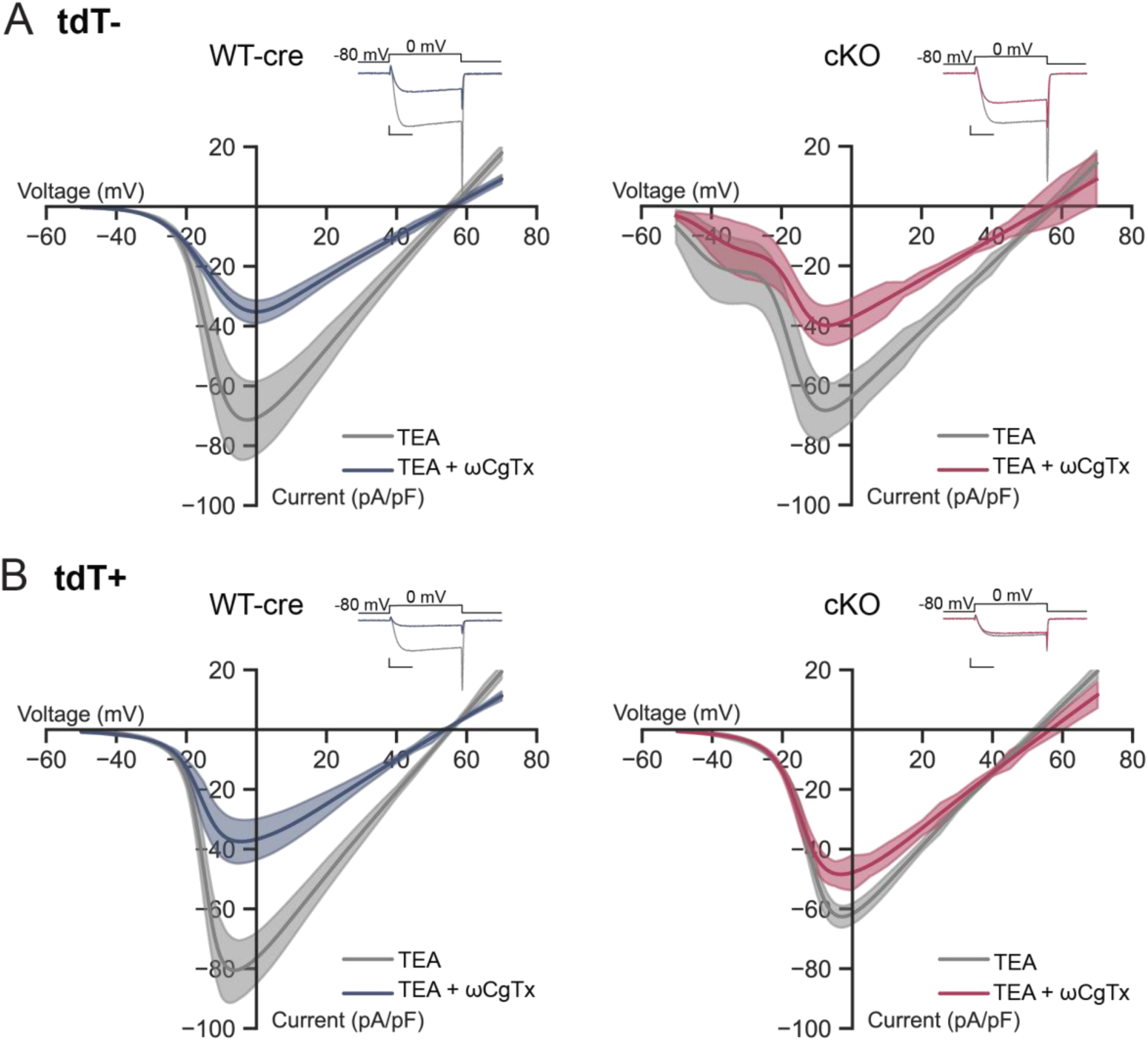
Whole cell calcium current recordings using ω-CgTx-GVIA confirms selective loss of Ca_V_2.2 channels in *Trpv1*-lineage neurons of *Cacna1b^fl/fl^/Trpv1^Cre/-^* mice. Whole cell Ca_V_ currents were recorded in *Trpv1*-negative (tdT−, *A*) and *Trpv1*-positive (tdT+, *B*) DRG neurons isolated from *Trpv1^Cre/Cre^* (WT-cre, *left*) and *Cacna1b^fl/fl^/Trpv1^Cre/-^* (cKO, *right*) mice. Ca_V_ currents were isolated using a TEA-based extracellular solution with 2 mM calcium as the charge carrier (gray) in the absence (gray) and presence (blue or red) of 2 µM ω-CgTx-GVIA to inhibit Ca_V_2.2 channels (*A, B*). Representative current recordings evoked by steps to 0 mV from -80 mV are shown in insets for each average current-voltage plot. Scale bars are 400 pA and 10 ms. *A,* Ca_V_ currents in tdT negative cells from WT-cre (*left, n* = 6) and in cKO (*right*, *n* = 5) mice before (gray) and after (blue/red) ω-CgTx-GVIA (tdT−: Control | ω-CgTx: *t* = -2.573, *p* = 0.028; Student’s t-test; tdT+: Control | ω-CgTx: *t* = -2.761, *p* = 0.025, Student’s t-test). *B*, Ca_V_ currents in tdT positive cells from WT-cre mice (n = 7, blue/*left*) were inhibited ∼50% by ω-CgTx-GVIA (*t* = -3.584, *p* = 0.005, Student’s t-test), whereas Ca_V_ currents in tdT positive cells from cKO mice (cKO, *n* = 7, red/*right*) were slightly, but not statistically smaller in the presence of ω-CgTx-GVIA (*t* = -1.893, *p* = 0.083; Student’s t-test). The reduced currents at the peak of the ω-CgTx-GVIA current-voltage relationship is likely due to rundown of Ca_V_ current which has been well documented particularly for Ca_V_1 channels (see Supplemental Fig. 1). Ca_V_ current-voltage relationships show average (solid line) ± SEM (shaded area) values together with the average fitted curves using one (*A, left; B*) or the sum of two (*A, right*) Boltzmann-linear line functions. Average parameters calculated from the fitted curves of individual cells.

### Selective inactivation of *Cacna1b* in *Trpv1*-lineage neurons is associated with increased neuronal excitability

After validating the cKO mouse model, we compared current clamp recordings from tdT+ WT-cre and cKO DRG neurons to assess the contribution of Ca_V_2.2 currents to overall excitability of *Trpv1*-lineage neurons. Average resting membrane potentials of tdT+/WT-cre (*n* = 14) and tdT+/cKO (*n* = 10) neurons were similar (WT-cre: -62 ± 2.58 mV; cKO: -63 ± 2.62 mV; *U* = 32.000, *p* = 0.4711; Mann-Whitney U), but neurons of cKO mice were more excitable; they fired action potentials in response to smaller currents (Fig. 4A, 4B; Rheobase: WT-cre = 1.09 ± 0.21 nA, cKO = 0.56 ± 0.12 nA; WT-cre | cKO: *t* = 2.160, *p* = 0.0435; Welch’s t-test) as well as more action potentials during the current step compared to WT-cre controls (Fig. 4A, 4B, 4D). The trajectory of action potentials in tdT+/WT-cre and tdT+/cKO neurons are compared in the dV/dt plots (Fig. 4C); action potentials in tdT+/cKO neurons exhibit steeper rising phases and deeper after hyperpolarizations compared to tdT+/WT-cre neurons (Fig. 4C). The threshold voltage at the inflection point of the first action potential generation (Fig. 4E; WT-cre = -33.93 ± 2.66 mV, cKO = -31.70 ± 2.61 mV; WT-cre | cKO: *t* = -0.579, *p* = 0.5683; Student’s t-test) and the average action potential amplitudes (Fig. 4F; WT-cre = 58.52 ± 3.75 mV, cKO = 65.04 ± 2.26 mV; WT-cre | cKO: *t* = -0.704, *p* = 0.4886; Student’s t-test) were the same in neurons of cKO and WT-cre mice. Action potential widths were shorter (Fig, 4G) in cKO neurons compared to WT-cre controls (WT-cre = 2.94 ± 0.22 ms, cKO = 2.45 ± 0.31 ms; WT-cre | cKO: *U* = 109.000, *p* = 0.0242; Mann-Whitney U).

**Figure 4.**
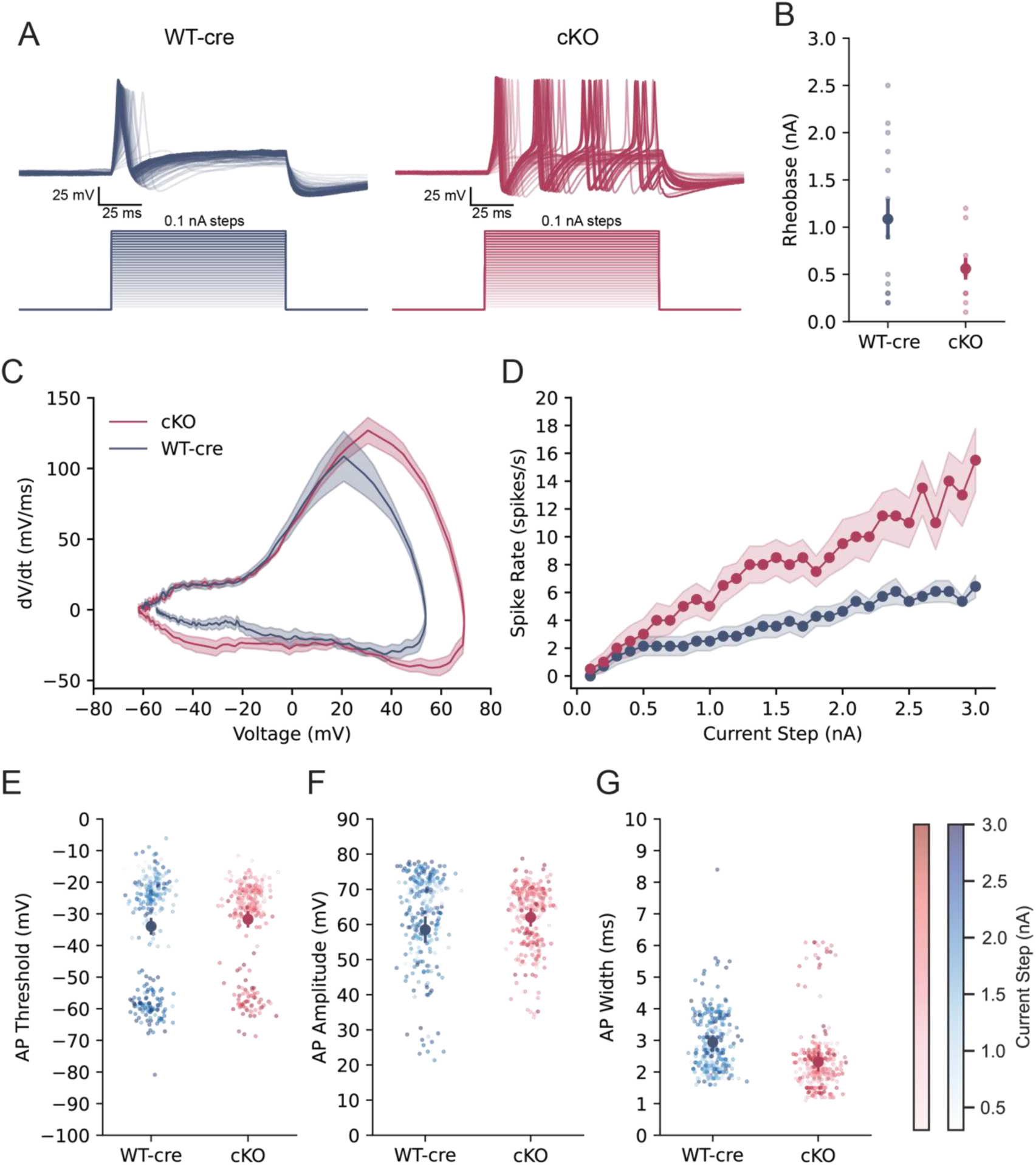
The absence of Ca_V_2.2 channels in *Trpv1*-lineage neurons is associated with increased cell excitability. Whole cell recordings from *Trpv1*-lineage DRG neurons (tdT+) were used to assess cell excitability in *Trpv1^Cre/Cre^* (WT-cre, dark blue; *n* = 14) and *Cacna1b^fl/fl^/Trpv1^Cre/-^* (cKO, red; *n* = 10) mice. Resting membrane potentials were not significantly different between WT-cre (-62.4 ± 2.6 mV) and cKO (-63.3 ± 2.6 mV) neurons (*U* = 32.0, *p* = 0.471; Mann-Whitney U). ***A***, Examples of voltage recordings from neurons of WT-cre (blue, *left*) and cKO (red, *right*) mice using sodium-based external solution with 2 mM calcium. Action potentials (APs) were evoked by current injections of 100 ms applied in 0.1 nA increments. In *Trpv1*-positive neurons from cKO mice, APs were triggered with smaller current injections (***B***), had slightly larger amplitudes (***A, C, F***) and occurred at higher frequencies (***D***), and had shorter widths (***C, G***) compared to WT-cre controls. Minimum currents required to evoke APs (Rheobase, ***B***) for WT-cre = 1.09 ± 0.21 nA, cKO = 0.56 ± 0.12 nA (WT-cre | cKO: *t* = 2.16, *p* = 0.0435; Welch’s t-test). Small symbols represent values from individual cells together with average ± SEM (solid circle, vertical bars) for each condition. ***C***, Average phase plots show dV/dt trajectory for first APs evoked by max current injection for WT-cre (*blue*) and cKO (*red*) recordings. Data shown are average ± SEM (solid line, shaded region). ***D***, Average AP firing rates as a function of current injection size. Current amplitude significantly affects spike rate (*β* = 4.413, *p* < 0.0001) and there is a significant interaction between mouse strain and current amplitude (WT-cre | cKO: *β* = -2.456, *p* < 0.0001; mixed-effects regression model). Values are average ± SEM (circles, shaded region). ***E, F, G,*** Features of action potentials evoked by current injections: ***E***, threshold voltage of first APs (WT-cre = -33.93 ± 2.66 mV, cKO = -31.70 ± 2.61 mV; WT-cre | cKO: *t* = -0.5793, *p* = 0.5683; Student’s t-test); ***F***, average AP amplitudes for each current injection (WT-cre = 58.52 ± 3.75 mV, cKO = 62.03 ± 2.64 mV; WT-cre | cKO: *t* = -0.7044, *p* = 0.4886; Student’s t-test); and ***G***, average AP widths for each current step (WT-cre = 2.94 ± 0.22 ms, cKO = 2.31 ± 0.303 ms; WT-cre | cKO: *U* = 109.000, *p* = 0.0242; Mann-Whitney U). Values are individual current injections that elicited an AP (small circles) together with mean (large circles) ± SEM (vertical line); current injection amplitudes are distinguished by the color gradient of the symbol (light to dark).

### Acute block of Ca_V_2.2 channels in *Trpv1*-lineage neurons has little effect on neuronal excitability

The increased neuronal excitability in *Trpv1*-lineage neurons of cKO mice compared to WT controls was unexpected. The faster rising phase of action potentials and increased firing rates are consistent with upregulation of Na_V_ channels which has been reported to occur following prolonged periods of neuronal inhibition^32,33^. We therefore tested the effects of acute inhibition of Ca_V_2.2 channels in *Trpv1*-lineage neurons using ω-CgTx-GVIA (Fig. 5, *n* = 11). Acute block of Ca_V_2.2 channels did not alter neuronal excitability (Fig. 6D; pre | post: *t* = -0.389, *p* = 0.7057; Paired t-test) and only slightly altered the action potential trajectory (Fig. 5A, 5B). First action potential thresholds (Fig. 5D; pre | post: *W* = 30.000, *p* = 0.8311; Wilcoxin signed-rank test), average action potential amplitude (Fig. 5E; pre | post: *t* = 0.974, *p* = 0.3530; Paired t-test), and average action potential width (Fig. 5F; pre | post: *W* = 16.000*, p* = 0.1475; Wilcoxin signed-rank test) were not significantly different before and after ω-CgTx-GVIA. The loss of Ca_V_2.2 channel function does not result in an increase in neuronal excitability and therefore suggests that the increase in neuronal excitability observed in *Trpv1*-lineage neurons from cKO mice may reflect a compensatory response to the loss of Ca_V_2.2 channel function and reduced synaptic transmission^10,32,33^.

**Figure 5.**
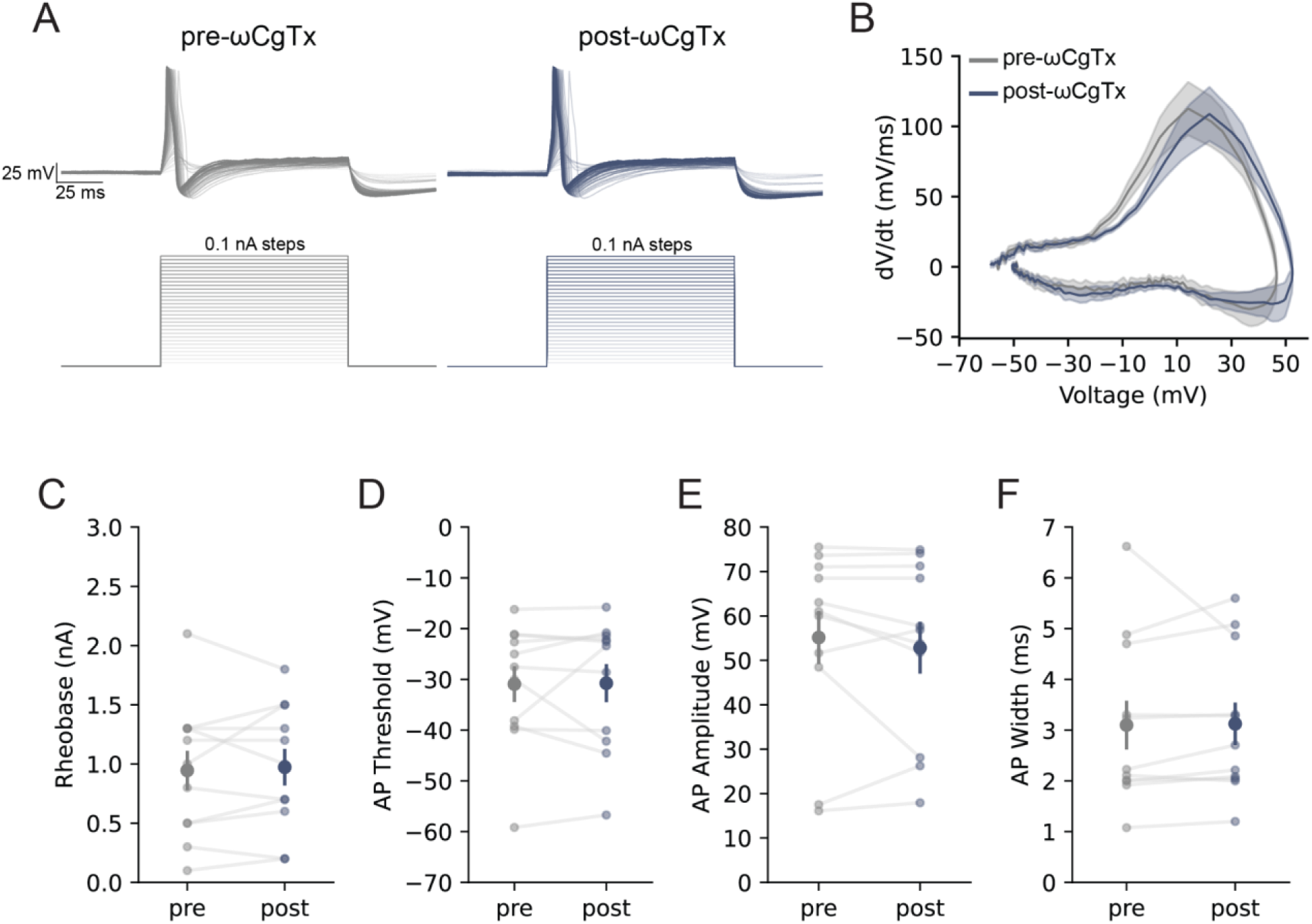
Acute pharmacological inhibition of Ca_V_2.2 currents in *Trpv1*-lineage neurons does not alter cell excitability. Whole cell voltage recordings from *Trpv1^Cre/Cre^* neurons (WT-cre, *n* = 11) in the absence (gray) and presence (dark blue) of ω-CgTx-GVIA. ***A***, Voltage recordings in the absence (gray, *left*) and presence (dark blue, *right*) of 2 µM ω-CgTx. Recordings were obtained using sodium-based external solution with 2 mM calcium. Action potentials were evoked by current injections of 100 ms applied in 0.1 nA increments. ***B***, average dV/dt phase plots of first action potentials evoked by max current injections in the absence (gray) and presence of ω-CgTx (dark blue). ***C***, ***D, E, F***, features of action potentials: amplitude of minimum current injected to trigger an action potential (***C***, Rheobase; before ω-CgTx = 0.95 ± 0.17 nA and after ω-CgTx = 0.97 ± 0.15 nA; pre | post: *t* = -0.3886, *p* = 0.7057; Paired t-test); ***D,*** first action potential thresholds (before ω-CgTx = -34.56 ± 3.73 mV; after ω-CgTx = -33.30 ± 3.78 mV; pre | post: *W* = 30.000, *p* = 0.8311; Wilcoxin signed-rank test); ***E***, average action potential amplitude for each current injection (before ω-CgTx = 55.15 ± 5.97 mV, after ω-CgTx = 52.84 ± 5.82 mV; pre | post: *t* = 0.9742, *p* = 0.3530; Paired t-test); and ***F***, average action potential widths for each current step (before ω-CgTx = 3.10 ± 0.48 ms, after ω-CgTx = 3.13 ± 0.42 ms; pre | post: *W* = 16.000, *p* = 0.1475; Wilcoxin signed-rank test). Values are individual cells (small circles) with mean (large circles) ± SEM (vertical lines). Horizontal lines connect individual cell recordings before and after ω-CgTx application.

**Figure 6.**
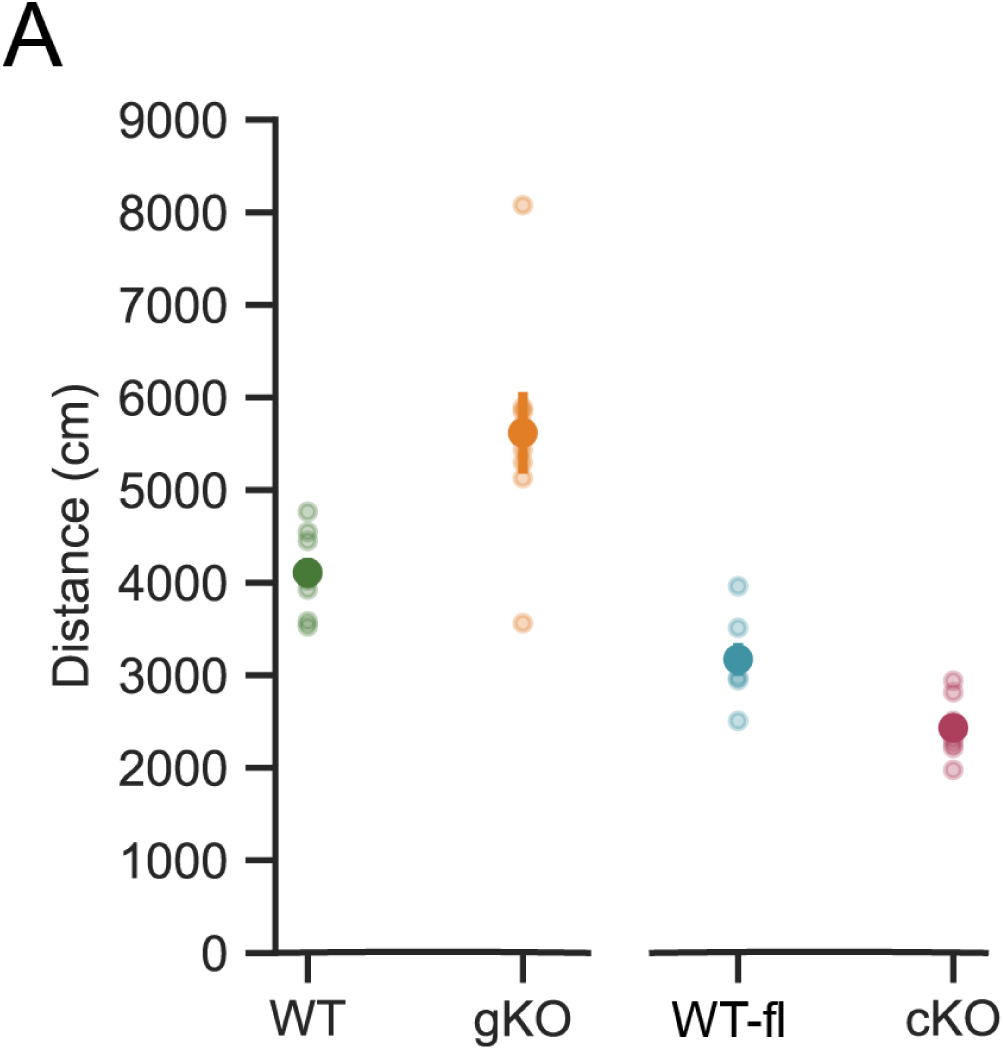
The hyperactivity behavior typical of global Ca_V_2.2 knockout mice is not a feature of conditional KO mice lacking Ca_V_2.2 in *Trpv1*-lineage neurons. Open field assay of total distance travelled (in 5 mins) for wildtype (WT, green; *n* = 8) and global *Cacna1b^-/-^* knockout (gKO, orange; n = 8); and for *Cacna1b^fl/fl^* controls (WT-fl, blue; *n* = 7) and conditional knockout *Cacna1b*^fl/fl^/*Trpv1^Cre/-^* (cKO, red; *n* = 7) mice were significantly different by strain (WT | gKO | WT-fl | cKO: *F* = 26.140, *p* < 0.0001; One-way ANOVA). WT = 4107.7 ± 158.2 cm, gKO = 5672.0 ± 438.9 cm, WT-fl = 3172.3 ± 174.5 cm and cKO = 2431.1 ± 130.1 cm (WT | gKO: *p* = 0.0009; WT-fl | cKO: *p* = 0.2619, One-way ANOVA, post hoc Tukey HSD). Values shown are measures from individual mice (small circles) together with mean (large darker circles) ± SEM (vertical lines).

### cKO and WT-fl control mice have similar overall activity levels

cKO and WT-fl mice exhibited similar levels of activity including distance travelled in an open field assay (Fig 6A; *F* = 26.140, *p* < 0.0001, One-way ANOVA; *p* = 0.2619 post hoc Tukey HSD). The WT-like behavior of cKO mice was very different from the pronounced hyperactivity of gKO mice^10,11^ (see supplemental video). In open field, gKO mice travelled 1.4 times the distance of WT controls (Fig. 6A; WT: *n* = 8, gKO: *n* = 8; WT | gKO: *p* = 0.0009; One-way ANOVA, post hoc Tukey HSD). Our data show that *Cacna1b* expression in non-*Trpv1* neurons underlies the pronounced hyperactivity phenotype associated with gKO mice^11^.

### Ca_V_2.2 channels in *Trpv1*-lineage neurons regulate paw withdrawal response thresholds to heat and cold

We measured paw withdrawal thresholds in WT-fl and cKO mice to heat, mechanical, and cold stimuli applied to the plantar surface of hind paws (Fig. 7A - 7C). *Cacna1b* is expressed in all sensory neurons^23,24^, but *Trpv1* has a more restricted expression pattern including in heat and cold responsive, but not mechanosensitive neurons^24,34^. In cKO mice, the delay to paw withdrawal following exposure to heat (Fig. 7A) and cold (Fig. 7B) stimuli were significantly longer compared to WT-fl controls (Fig. 7A, 7B; Heat: *U* = 20.000, *p* = 0.0001; Cold: *U* = 33.000, *p* = 0.0444; Mann-Whitney U). Whereas stimulus-response thresholds to mechanical stimuli were not statistically different between cKO and WT-fl mice (Fig. 7C; *t* = 1.311, *p* = 0.2002; Welch’s t-test). These results illustrate cell-specific deletion of *Cacna1b* in cKO mice, confirm the involvement of Ca_V_2.2 channels in withdrawal responses to heat^4,5,10^ and demonstrate a role for Ca_V_2.2 channels in regulating response thresholds to cold (Fig. 7B).

**Figure 7.**
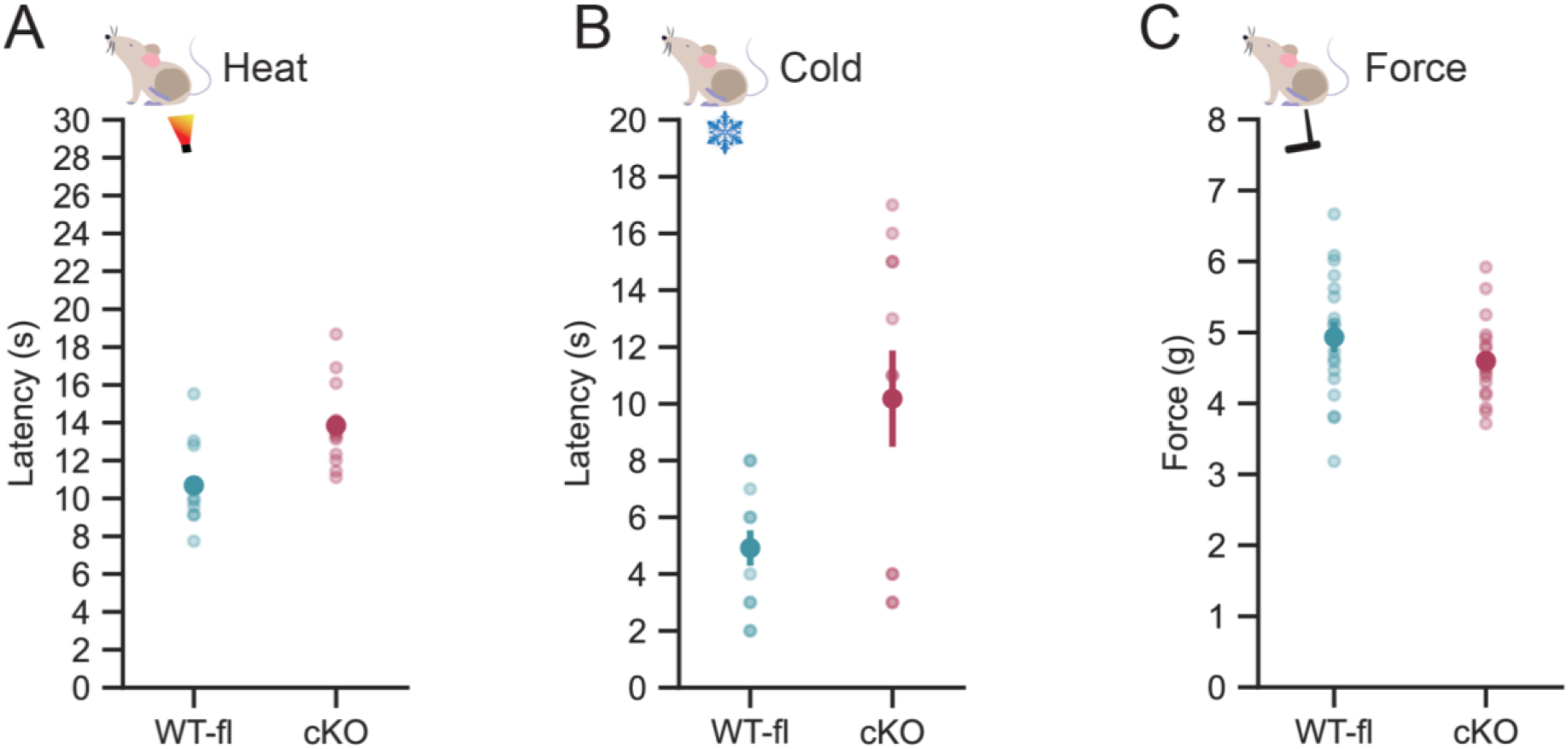
Reduced paw withdrawal sensitivity to heat and cold stimuli reduced in mice lacking Ca_V_2.2 channels in *Trpv1*-lineage neurons compared to wild-type. Paw withdrawal responses to heat (*A*, latency), cold (*B*, latency) and force (*C*, grams) were measured in *Cacna1b*^fl/fl^ (WT-fl, blue symbols) and *Cacna1b^fl/fl^/Trpv1^Cre/-^* (cKO, red symbols) mice (WT-fl are littermate controls). Responses to heat (*A*; *n* = 15) and force (*C*; *n* = 18) are the baseline responses for both capsaicin and saline injected mice used in Fig. 8. Small symbols represent averages of 6 measurements from each mouse (3 from each hindpaw) together with mean values (large symbols) ± SEM (vertical bars) for each genotype. Cold responses (*B*) were measured from a separate set of cKO (*n* = 11) and WT-fl (*n* =12) littermate control mice. Comparison of responses to heat (*A)* for WT-fl = 10.69 ± 0.48 s and cKO = 13.85 ± 0.53 s (WT-fl | cKO: *U* = 20.000, *p* = 0.0001; Mann-Whitney U); cold (*B*) for WT-fl = 4.92 ± 0.62 s and cKO = 10.18 ± 1.69 s (WT-fl | cKO: *U* = 33.0000, *p* = 0.0444; Mann-Whitney U); and force (*C*) for WT-fl = 4.93 ± 0.22 g and cKO = 4.60 ± 0.14 g (WT-fl | cKO: *t* = 1.2109 , *p* = 0.2002; Welch’s t-test).

### Ca_V_2.2 channels in *Trpv1*-lineage neurons necessary for capsaicin-induced rapid heat, but not mechanical hypersensitivity

We previously reported that Ca_V_2.2 channels were essential for the development of capsaicin-induced heat, but not mechanical hypersensitivity in skin, using a gKO mouse model and local intradermal injection of ω-CgTx-MVIIA in wild-type mice^4,10^. Our data suggested that Ca_V_2.2 channels in *Trpv1* neurons were critical for capsaicin-induced hypersensitivity, but we could not rule out involvement of non-*Trpv1* expressing cells^4,10^. We therefore compared withdrawal thresholds in cKO and WT-fl littermate controls to heat (Hargreaves, Fig. 8A) and mechanical (von Frey, Fig 8B) stimuli before, 15 min, and 30 min following intradermal administration of 0.1% w/v capsaicin or saline.

**Figure 8.**
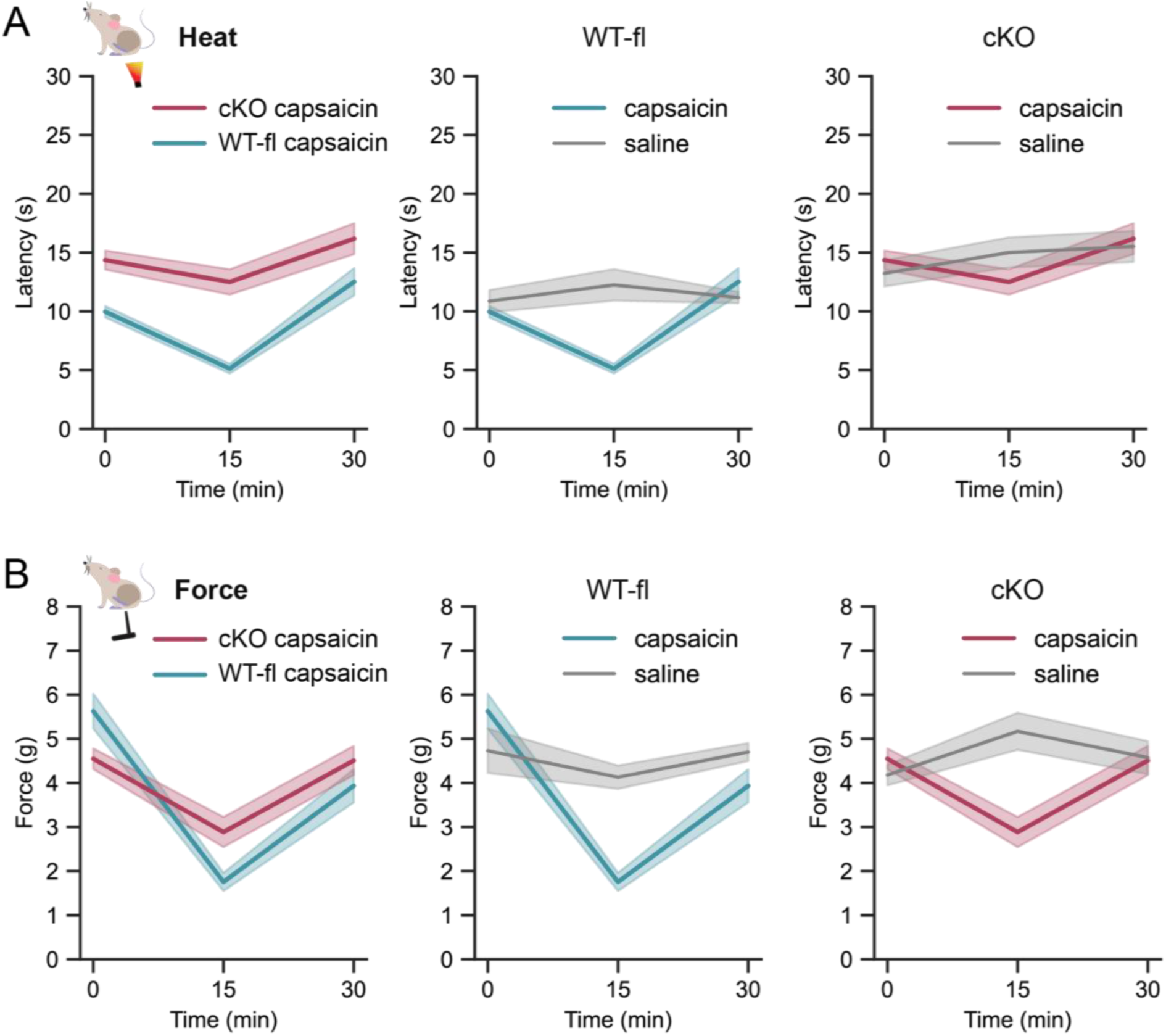
Capsaicin-induced heat hypersensitivity absent in mice lacking Ca_V_2.2 channels in *Trpv1*-lineage neurons. Paw withdrawal responses measured at 0, 15 min and 30 min intervals from wild-type *Cacna1b*^fl/fl^ (WT-fl, blue; ***A****, **B***) and *Cacna1b^fl/fl^/Trpv1^Cre/-^* (cKO, red; ***A****, **B***) mice to radiant heat (latency, ***A***) and force (threshold, ***B***) applied to plantar hind paw. Right paws: *i.d.* 20 μl, 0.1% capsaicin (blue, red) or 20 μl, 0.1% saline (gray). Values shown are mean (solid line) ± SEM (shaded area). ***A***, Saline (*n* = 7, gray) and capsaicin (*n* = 8, blue). At 15 min, WT-fl mice (*left*): saline = 12.25 ± 1.34 s, capsaicin = 5.13 ± 0.41 s (saline | capsaicin: *F* = 20.580, *p* = < 0.0001, Two-way ANOVA; *p* = 0.0004 post hoc Tukey HSD). cKO mice (*right*): saline = 15.00 ± 1.28 s, capsaicin = 12.50 ± 1.08 s (saline | capsaicin: *F* = 20.580, *p* = < 0.0001, Two-way ANOVA; *p* = 0.3600, post hoc Tukey HSD). ***B***, Saline (*n* = 7, gray) and capsaicin (*n* = 11, blue). At 15 min, WT-fl mice (*left*): saline = 4.13 ± 0.26 g, capsaicin = 1.75 ± 0.20 g (saline | capsaicin: *F* = 53.310, *p* < 0.0001, Two-way ANOVA; *p* < 0.0001, post hoc Tukey HSD). cKO mice (*right*) saline (*n* = 9, gray) and capsaicin (*n* = 9, red). Saline = 4.13 ± 0.26 g, capsaicin = 2.89 ± 0.34 g (saline | capsaicin: *F* = 53.310, *p* < 0.0001, Two-way ANOVA; *p* < 0.0001, post hoc Tukey HSD).

The capsaicin-induced hypersensitivity response in skin develops within minutes and is robust at 15 mins post capsaicin injection^4,10^. At this time point, WT-fl mice exhibited capsaicin-induced hypersensitivity to both heat (Fig. 8A, *middle*; saline: *n* = 7, capsaicin: *n* = 8; saline | capsaicin: *F* = 20.580, *p* = 0.0001, Two-way ANOVA; *p* = 0.0004, post hoc Tukey HSD) and mechanical (Fig. 8B, *middle*; saline: *n* = 7, capsaicin: *n* = 11; saline | capsaicin: *F* = 53.310, *p* < 0.0001, Two-way ANOVA; *p* = 0.0001, post hoc Tukey HSD) stimuli compared to saline controls. By contrast, cKO mice failed to develop capsaicin-induced hypersensitivity to heat; average paw withdrawal latencies for capsaicin-injected and saline-injected paws were not different (Fig. 8A, *right*; capsaicin: *n* = 8, saline: *n* = 7) (saline | capsaicin: *p* = 0.3600; Two-way ANOVA, post hoc Tukey HSD). Whereas capsaicin-induced mechanical hypersensitivity was robust and significantly different from saline control responses (Fig. 8B, *right*; mechanical stimuli, capsaicin: *n* = 9, saline: *n* = 9; saline | capsaicin: *p* = 0.0001; Two-way ANOVA, post hoc Tukey HSD). These results show that Ca_V_2.2 channels expressed in *Trpv1*-lineage neurons are the source of depolarization-induced calcium entry and this event is essential for the development of rapid, short-term hypersensitivity to heat.

### Ca_V_2.2 channels in *Trpv1*-lineage neurons contribute to the prolonged time course of CFA-induced heat, but not mechanical hypersensitivity

We previously reported that Ca_V_2.2 channels are essential for heat hypersensitivity associated with chronic inflammation following intraplantar CFA^5^. CFA-induced heat hypersensitivity in response to intradermal CFA in gKO mice was eliminated^5^. Here we use the cKO mouse model to measure paw withdrawal responses to heat (Hargreaves, Fig. 9A) and mechanical (von Frey, Fig. 9B) stimuli before and for 7 days following intraplantar administration of CFA to assess the involvement of Ca_V_2.2 channels expressed in *Trpv1*-lineage neurons. In WT-fl mice (*n* = 8), CFA-induced heat (ipsi | contra: day 1, *p* < 0.0001; day 2, *p* < 0.0001; day 3, *p* < 0.0001; day 4, *p* = 0.0084; day 5, *p* = 0.0042; day 6, *p* = 0.008; day 7, *p* = 0.0007; repeated measures ANOVA, post hoc Tukey HSD) and mechanical hypersensitivity (ipsi | contra: day 1, *p* = 0.0001; day 2, *p* < 0.0001; day 3, *p* < 0.0001; day 4, *p* < 0.0001; day 5, *p* = 0.0003; day 6, *p* = 0.0003; day 7, *p* = 0.0001; repeated measures ANOVA, post hoc Tukey HSD) was different in ipsilateral, compared to contralateral paws, for all 7 days of the measurement period. In cKO (*n* = 8) mice, CFA-induced heat hypersensitivity was different from day 1 to 3, but on days 4-7 the response was curtailed and responses in contralateral and ipsilateral paws were not significantly different (ipsi | contra: day 1, *p* = 0.0079; day 2, *p* = 0.002; day 3, *p* = 0.0002; day 4, *p* = 0.3286; day 5, *p* = 0.1545; day 6, *p* = 1766; day 7, *p* = 0.7123). By contrast, mechanical hypersensitivity was different in ipsilateral compared to contralateral paws for all 7 days (ipsi | contra: day 1, *p* = 0.0002; day 2, *p* < 0.0001; day 3, *p* < 0.0001; day 4, *p* = 0.0017; day 5, *p* = 0.0015; day 6, *p* = 0.0002; day 7, *p* = 0.0067; repeated measures ANOVA, post hoc Tukey HSD). Mechanical hypersensitivity following CFA administration remained intact in the absence of Ca_V_2.2 channels on *Trpv1*-lineage neurons.

**Figure 9.**
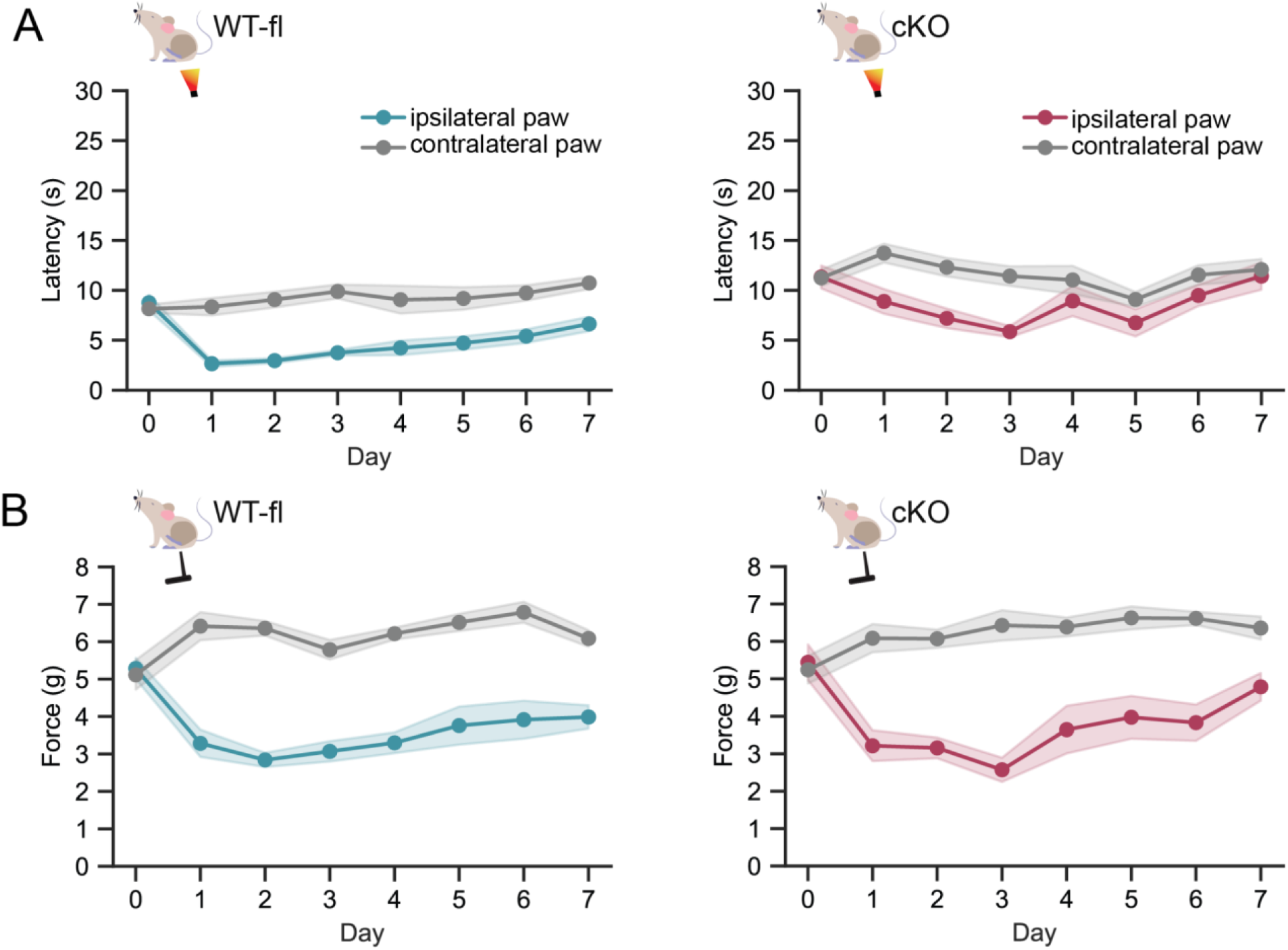
Ca_V_2.2 channels in *Trpv1*-lineage neurons contribute to the late phase of CFA- induced heat hypersensitivity. Paw withdrawal responses to heat (A, latency) and mechanical (B, force) stimuli applied to the plantar surface of mouse hind paws. Contralateral (gray) and ipsilateral (blue and red) paw responses measured daily from wild-type *Cacna1b*^fl/fl^ (WT-fl, blue/left; ***A****, **B***) and *Cacna1b*^fl/fl^/*Trpv1^Cre/-^* (cKO, red/right; ***A****, **B***) mice before, and for 7 days after ipsilateral paw injected with *i.d.* 20 μl CFA. Values shown are mean (solid line) ± SEM (shaded area). ***A***, Heat WT-fl (*n* = 8, *left*) main effect by day (ipsi | contra: *F* = 3.20, *p* = 0.0072), injection (ipsi | contra: *F* = 181.94, *p* < 0.0001), and day by injection interaction (ipsi | contra: *F* = 8.44, *p* < 0.0001; repeated measures ANOVA). For each of day 1-7 days post-CFA (ipsi | contra: day 1, *p* < 0.0001; day 2, *p* < 0.0001; day 3, *p* < 0.0001; day 4, *p* = 0.0084; day 5, *p* = 0.0042; day 6, *p* = 0.008; day 7, *p* = 0.0007; repeated measures ANOVA, post hoc Tukey HSD). cKO (*n* = 8, *right*) main effect by day (ipsi | contra: *F* = 2.30, *p* = 0.0412), injection (ipsi | contra: *F* = 121.84, *p* < 0.0001), and day by injection interaction (ipsi | contra: *F* = 3.71, *p* = 0.0027; repeated measures ANOVA). cKO 1-7 days post-CFA (ipsi | contra: day 1, *p* = 0.0079; day 2, *p* = 0.002; day 3, *p* = 0.0002; day 4, *p* = 0.3286; day 5, *p* = 0.1545; day 6, *p* = 1766; day 7, *p* = 0.7123; repeated measures ANOVA, post hoc Tukey HSD). ***B*,** Mechanical responses: WT-f (*n* = 7, *left*) main effect by day (ipsi | contra: *F* = 2.33, *p* = 0.0420), injection (ipsi | contra: *F* = 133.74, *p* < 0.0001), and day by injection interaction (ipsi | contra: *F* = 14.63, *p* < 0.0001; repeated measures ANOVA). Ipsilateral and contralateral responses compared days 1-7 post-CFA (ipsi | contra: day 1, *p* = 0.0001; day 2, *p* < 0.0001; day 3, *p* < 0.0001; day 4, *p* < 0.0001; day 5, *p* = 0.0003; day 6, *p* = 0.0003; day 7, *p* = 0.0001; repeated measures ANOVA, post hoc Tukey HSD). cKO (*n* = 7, *right*) main effect by day (ipsi | contra: *F* = 3.57, *p* = 0.0042), injection (ipsi | contra: *F* = 37.37, *p* = 0.0009), and day by injection interaction (ipsi | contra: *F* = 6.04, *p* = 0.0001; repeated measures ANOVA). cKO ipsilateral and contralateral responses compared days 1-7 post-CFA (ipsi | contra: day 1, *p* = 0.0002; day 2, *p* < 0.0001; day 3, *p* < 0.0001; day 4, *p* = 0.0017; day 5, *p* = 0.0015; day 6, *p* = 0.0002; day 7, *p* = 0.0067; repeated measures ANOVA, post hoc Tukey HSD).

These data, combined with our previous studies of gKO mice which do not exhibit CFA-induced heat hypersensitivity^5^, suggest that Ca_V_2.2 channels in both *Trpv1*-lineage neurons and *Trpv1*- lineage cells combined are required for heat hypersensitivity following chronic neuroinflammation induced by CFA.

## DISCUSSION

Voltage-gated Ca_V_2.2 channels control presynaptic calcium entry and neurotransmitter release at many synapses^1,2^. Studies using both global knockout of Ca_V_2.2 channels and local inhibition through small molecule blockers have also established a critical role for Ca_V_2.2 channels in the transmission of pain information^4,5,10,35,36^. Furthermore, a peptide inhibitor of Ca_V_2.2 channels administered intrathecally is analgesic in patients with otherwise intractable pain^37,38^. More recently, our lab reported new data showing a critical and specific role of Ca_V_2.2 channels at peripheral sites in skin, in the development of heat but not mechanical hypersensitivity^4,5,10^. Ca_V_2.2 channels are enriched in *Trpv1*-expressing, heat-sensitive neurons^24^, so we used a *Trpv1*- conditional gene inactivation strategy to: 1) establish the relative contribution of Ca_V_2.2 channels in *Trpv1-*expressing and *Trpv1*-non-expressing neurons to behavioral responses associated with neuroinflammatory events, and 2) establish the acute and longer term consequences of eliminating the activity of Ca_V_2.2 channels on neuronal excitability. In our studies we also validated the Cre-dependent mouse model for use to explore and define the contributions of the *Cacna1b* gene and Ca_V_2.2 channels in other cell types for *in vivo*, circuit and cell-based studies.

Although not ideal for *in vivo* studies, because of the pronounced hyperactivity phenotype, global *Cacna1b* KO mice proved critical as a negative control for Ca_V_2.2 protein analysis. Specifically, to identify Ca_V_2.2 specific bands in native tissue, and to reveal tissue-specific difference in protein expression. The polyclonal anti-Ca_V_2.2 used here detect Ca_V_2.2-specific proteins and they have been used by our lab^10,18,19^ but they also binds several non-Ca_V_2.2 proteins in native tissue which can only be identified with certainty using an appropriate negative control. The non-Ca_V_2.2 bands labelled by polyclonal antibodies raised against Ca_V_2.2 have typically not been shown in published data^20,39–42^ but, as we show here and in our previous publications^10,18–20^, non-Ca_V_2.2 signals are a prominent feature in Western analyses of native tissues. By comparing to gKO protein preparations, we also demonstrated tissue-specific differences in the pattern of both non-specific signals (e.g. a non-Ca_V_2.2 signal at 125 kDa in brain is absent in DRG; Fig 1B) as well as Ca_V_2.2-specific protein signals (an additional ∼240 kDa Ca_V_2.2-specific protein band in DRG but not in brain; Fig. 1B)^20^, which are easily overlooked without comparing to a gKO negative control.

The heterogeneity of neurons in DRG can make it difficult to identify cell-specific differences in the biophysical properties of specific classes of neurons. We therefore combined the use of Cre-reporter mice for gene-specific inactivation of *Cacna1b* with AAV viral delivery of a Cre-dependent tdTomato (tdT) reporter to separate neurons visually into Cre- and non-Cre-expressing cells (Cre-positive/tdT+ and Cre-negative/ tdT−) for biophysical analyses of individual cells. We used both voltage and current clamp recordings to establish specificity of the cKO model, and we showed that the trajectory of action potentials in Cre-expressing neurons from cKO animals were slightly shorter from WT-cre control neurons (Fig. 4C, 4G). A subset of nociceptor neurons have a characteristic plateau phase which has been partly attributed to the presence and action of calcium channels^43–45^, and our data show that the percentage of cells with longer half widths is reduced in recordings from cKO mice, supporting a role for Ca_V_2.2 channels in defining the action potential plateau in *Trpv1*-expressing neurons (Fig. 4G).

We also assessed the impact of Ca_V_2.2 channels on the overall excitability of *Trpv1*-lineage neurons and unexpectedly found that *Trpv1*-lineage neurons from cKO mice exhibited a gain of function phenotype (Figs. 2-4). *Trpv1*-lineage neurons from cKO mice were more excitable as compared to WT-cre controls based on several criteria including, the amount of current required to fire an AP and the number of APs during a current step (Fig. 4). The biophysical features of *Trpv1*-lineage neurons in cKO mice were unexpected based on the behavioral phenotype of cKO mice, which were overall *less* sensitive to sensory stimuli (Figs. 7-9). We conclude that there is a compensatory increase in membrane excitability in cKO neurons associated with the absence of Ca_V_2.2 channel activity, most likely because of the reduction in the efficacy of synaptic transmission^10^. Data supporting the presence of a compensatory change in neuronal excitably includes showing that acute inhibition of Ca_V_2.2 channel activity in *Trpv1*-lineage neurons of WT-cre mice does not alter neuronal excitability and only slightly reduced the action potential half width (Fig. 5). Additionally, analyses of cell excitability and action potential trajectories are consistent with the upregulation of voltage-gated Na+ currents, as seen by a reduced current required to trigger AP generation, no difference in the resting membrane potential, and a more depolarized peak value of action potentials in *Trpv1*-lineage neurons of cKO mice (Fig. 4).

Compensatory homeostatic mechanisms in neurons have been reported following prolonged changes in synaptic efficacy^32,33,46^. In tectal neurons, changes in the level of Na_V_ channels underlies the homeostatic mechanisms used to maintain a constant input–output function during development^47^, and the expression of Na_V_ channels and other ion channels changes in models of multiple sclerosis to compensate for reduced signal conductance following demyelination^48–50^.

In our studies, the increase in neuronal excitability of *Trpv1*-neurons measured at the single cell level in cKO mice compared to WT-cre controls did not fully compensate for change in synaptic strength. *Trpv1*-cKO mice have reduced responsiveness to heat and cold stimuli, and gKO mice are also less sensitive to mechanical stimuli^4,5,10^. Furthermore, *Cacna1b* cKO and gKO mice do not develop heat hypersensitivity in response to capsaicin and have shorter (cKO) or no heat hypersensitivity (gKO) response to CFA suggesting that the loss of Ca_V_2.2 channels dominates in defining the neuroinflammatory response in skin^4,5,10^. Thus, neuronal excitability can change quite substantially in *Trpv1*-expressing neurons in response to prolonged alterations in synaptic transmission without impacting overall behavior. Our observations might have relevance to current therapeutic strategies aimed at reducing neuronal excitability (e.g. blocking Na_V_1.8 and Na_V_1.7) for pain management^51–55^. Sensory neurons undergo dynamic and, as we show here, remarkably sustained changes in excitability so while it is possible that targeting ion channels to down regulate neuronal excitability could be effective for short-term analgesia, an inhibitor of transmitter release from peripheral nerve endings and/or presynaptic termini in the spinal cord might prove more effective for longer term, chronic pain management.

The use of the cKO approach reported here, allowed us to define cell-specific actions of Ca_V_2.2 channels in mediating in heat hypersensitivity using both acute (capsaicin, Fig. 8) and long-lasting (CFA, Fig. 9) models of neuroinflammation. We reenforced and extended our previous findings using *Cacna1b* gKO and pharmacological approaches^4,5,10^ to show that Ca_V_2.2 channel activity in *Trpv1*-lineage neurons is the key initiating signal mediating capsaicin-induced heat hypersensitivity, while having no effect on the development of mechanical hypersensitivity in the same animals. Other signaling molecules independent of Ca_V_2.2 channel activity must mediate mechanical hypersensitivity. We know from our previous studies that IL-1a is a key player in capsaicin-induced heat and mechanical hypersensitivity^4^ so it is possible that capsaicin acts on TRPV1 receptors on immune cells^56–59^ to trigger cytokine release with the potential to sensitize mechanosensitive neuron populations.

The importance of non-*Trpv1*-expressing cells in mediating long-lasting inflammation triggered by intraplantar CFA is shown conclusively in our studies using cKO mice (Fig. 9). Whereas CFA failed to induced heat hypersensitivity in gKO mice^5^, in cKO mice heat hypersensitivity developed but the duration of the response was shortened. CFA-induced mechanical hypersensitivity in cKO mice was unaffected by the absence of Ca_V_2.2 channels. It is likely that calcium influx through Ca_V_2.2 channels plays a critical role in triggering the release of pro-inflammatory mediators, engaging immune cells and initiating an inflammatory cascades that promote pain and inflammation^4,5,36^. Inhibiting Ca_V_2.2 channels—and thereby disrupting these cascades—may help promote recovery in models of neuroinflammation.

Here, we provide conclusive evidence that Ca_V_2.2 channels expressed in *Trpv1*-lineage neurons are critical for the development of heat hypersensitivity. Ca_V_2.2 channels are the critical link between prolonged membrane depolarization of *Trpv1* neurons (e.g. by capsaicin) to neuroinflammation that results in a rapid increase in the responsiveness of sensory nerve endings to heat. Our studies also uncover a significant change in the excitability of sensory neurons in response to the loss of Ca_V_2.2 channels. Interestingly, this compensatory increase in excitability of *Trpv1*-expressing neurons is not manifested at the level of behavior which is dominated by Ca_V_2.2-dependent signaling at peripheral and presynaptic sensory nerve endings.

Given the dynamic changes in neuronal excitability observed here using the *Cacna1b* cKO model and the known developmental changes in *Trpv1* expression^60,61^, it will be important to assess the effects of cell-specific and temporally controlled inactivation of *Cacna1b* during development.

## MATERIALS AND METHODS

### Experimental animals

All mice used were bred at Brown University, and all protocols and procedures were approved by the Brown University Institutional Animal Care and Use Committee (IACUC). Mice were housed in a 12 hr light/dark cycle and tested between 10 am and 6 pm during the light cycle. 3 - 6-month-old male and female mice were used in all experiments. Experimenters were blind to animal genotype and solution injected for the experimental and data analysis steps.

*Cacna1b*^fl/fl^ mice were generated by Biocytogen in the C57BL6/J background using the CRISPR/Cas9 Extreme Genome Editing (CRISPR/EGE^TM^) platform. To enable Cre-mediated recombination, loxP sites were inserted in intron 4-5 and intron 5-6, both oriented in the 5’ direction. Cre-mediated recombination deletes constitutive exons 5 and 6 resulting in a frame shift and premature stop codon. The sequences of guide RNAs (sgRNA) used in CRISPR/EGT^TM^ and of primers for the targeting vector are in Table 2. The 5.6 kb targeting vector contained 1.5 kb homologous 5’ and 3’ arms.

*Cacna1b^f^*^l/fl^ were crossed to *Cacna1b^fl/fl^/Trpv1^Cre/-^* mice to generate F1 progeny (∼50% *Trpv1*- lineage specific *Cacna1b* knockout (cKO) and ∼50% *Cacna1b^f^*^l/fl^ littermate wild-type controls. Routine genotyping was performed by qPCR from ear punch biopsies. Probes were designed to detect the 5’ loxP site between exons 5 and 6, the *Cacna1b* WT allele, and *Cacna1b* allele with the intended deletion. We vigilantly monitored for the presence of mosaic mice for the *Cacna1b* deletion in our crosses. Mosaic mice were not used for experiments or breeding.

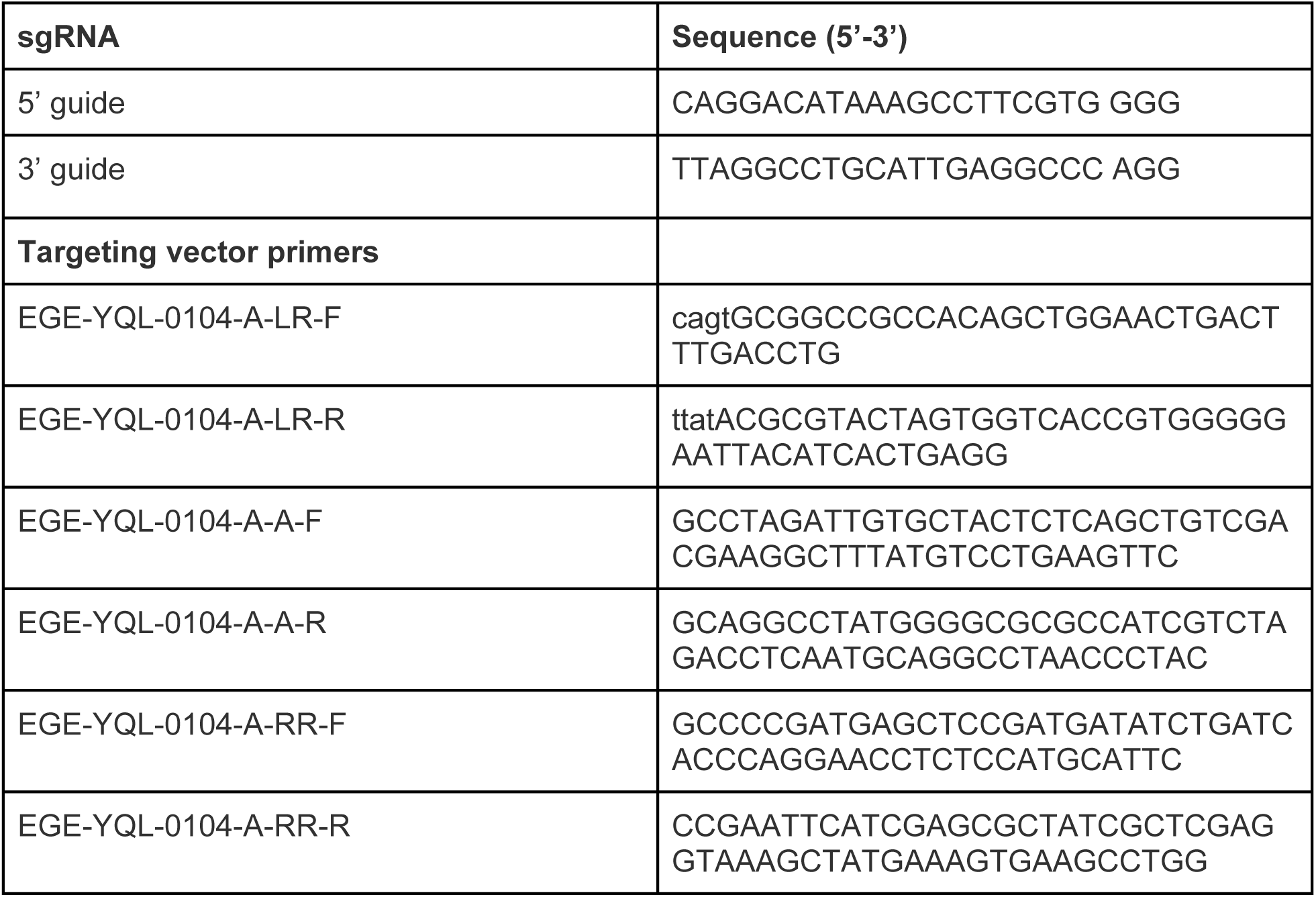

The *Cacna1b^−/−^* global deletion (gKO) mouse strain (*Cacna1b*tm5.1DiLi, MGI) generated in our lab has been described previously ^10^. A wild-type strain was bred in parallel from the same genetic background and used for comparison in hyperactivity experiments.

### Western blot analysis

Harvested brain was immediately homogenized on ice in Syn-PER buffer (87793, Thermo Fisher) containing protease inhibitor (Complete, mini; Roche Diagnostics). The homogenate was centrifuged at 1200 xg for 10 min at 4°C, supernatant transferred to a clean tube and centrifuged at 15000 xg for 30 min at 4°C and the pellet resuspended in ice-cold CL-91 buffer (CL-91–01, Logopharm) containing a protease inhibitor (Halt, Thermo Fisher). DRG were pooled from 5 mice, homogenized on ice in RIPA buffer (25 mM Tris-HCl, 150 mM NaCl, 1% NP-40, 1% sodium deoxycholate, 0.1% SDS pH 7.6) containing protease inhibitor (Complete, mini; Roche Diagnostics). After 3 hr incubation on ice, the homogenate was centrifuged at 10000 xg for 10 min and the supernatant was collected. Homogenate protein concentrations were estimated using BCA Protein Assay and BSA as a standard (23225, Pierce, Thermo Scientific).

Protein preparations were separated in tris-acetate 3-8% polyacrylamide gels (EA03752, Nu Page Thermo Fisher). Protein samples were incubated at room temperature for 20 min with Laemmli protein sample buffer (200 mM Tris-HCl, 2% w/v SDS, 10% v/v Glycerol, 100 mM DTT, pH 6.8). 20 µg of protein were loaded per well and the electrophoresis was run at 140 V (constant) using in MES-SDS as a running buffer (50 mM MES, 50 mM Tris Base, 0.1% SDS, 1 mM EDTA, pH 7.3, Thermo Fisher). After electrophoresis, proteins were transferred from polyacrylamide gels to 0.2 µm PVDF membranes using a semi dry transference system (Biorad Trans-Blot Turbo) at 1.3 A, 25 V for 10 min with a transfer buffer (Trans-Blot Turbo 5X, 20% v/v ethanol, Bio-Rad). Total protein was estimated using the LI-COR Revert™ 700 Total Protein Stain, and imaging was performed on a Bio-Rad ChemiDoc MP Imaging System. Total protein was visualized at IRDye 680.

Transfer membranes were blocked with 5% skimmed milk in PBS-Tween for 1 hr (PBST, 137 mM NaCl, 2.7 mM KCl, 10 mM Na_2_HPO_4_, 2 mM KH_2_PO_4_, 0.1% v/v Tween-20). Membranes were incubated with polyclonal rabbit anti-Ca_V_2.2 (ACC-002, Alomone Labs; 1:200 dilution in PBST, 2% w/v BSA) for 1 hr at room temperature, and detected using an anti-rabbit IgG (whole molecule) peroxidase conjugated antibody (Jackson Inmunoresearch 711-036-152; 1:10000 dilution in PBST, 2 % w/v BSA, 1h, room temp). Membranes were washed with PBST between each step in the protocol. Immunoblot membranes were processed using the SuperSignal West Pico PLUS Chemiluminescent substrate and the image was acquired using a Chemidoc MP (Bio-Rad).

Images were analyzed using Image Lab (version 6.1.0 build 7, Bio-rad). Band intensities were calculated and normalized to total protein for each lane. Protein intensities were calculated by normalizing to the control group. Adjustments of image brightness and contrast was applied to the entire gel.

### Cell isolation and electrophysiology

To visually identify *Trpv1*-lineage neurons for recording we used a tdTomato reporter AAV viral vector. Cre-dependent tdTomato expression was achieved by intrathecal injection of 10 µL of pAAV-FLEX-tdTomato packaged with AAV-PHP.s (AAV-PHP.s-FLEX-tdTomato; Addgene #: 28306-PHP.S, titer: 2E13 GC/mL) delivered at least 2 weeks prior to DRG isolation. For intrathecal injections, mice were lightly anesthetized with 2% isoflurane. A 26-gauge needle was manually inserted between L5 and L6 vertebrae and solution was injected once a tail flick was evoked (method adapted from Li *et al*.)^62^. A WPI microinjector (Model UMP3 Serial 191648 M06B; Sarasota, Florida USA) was used to inject virus at a flow rate of 250 nL/s. Mice were monitored for any abnormalities in gait after recovery.

Bilateral L3-L5 DRG were isolated 2-3 weeks post-injection from 8-10-week-old isoflurane euthanized mice. DRG were kept on ice in Hank’s Balanced Salt Solution (HBSS, GIBCO) before being transferred to HBSS containing 2 mg/mL collagenase (GIBCO) and 2.5 mg/mL trypsin (Sigma) at 37°C for 30 min. The pellet was lightly disturbed every 10 min. Protease activity was halted with the addition of Dulbecco’s Modified Eagle Medium (DMEM, GIBCO) supplemented with 10% fetal bovine serum (v/v, FBS, GIBCO) and 1 ng/mL nerve growth factor (NGF, Sigma). Single cells were isolated by trituration using a fire polished Pasteur pipette and resuspended in 500 µL 10% FBS-DMEM supplemented with 1 ng/mL NGF. Neurons were kept in suspension at 37°C with 5% CO_2_ until needed. 40 µL of the cell suspension was added to the center of the recording chamber and, after a 20 min settling time, the recording chamber was flooded with the Na-based external recording solution.

The external Na-based solution contained 135 mM NaCl, 2 mM CaCl_2_, 10 mM HEPES, 4 mM MgCl_2_, 2 mM KCl, adjusted to a pH of 7.4 with NaOH and to an osmolarity of 305 mOsm with NaCl; the TEA-based external solution used to isolate voltage-gated Ca^2+^ currents contained 135 mM TEA-Cl, 2 mM CaCl_2_, 10 mM HEPES, 4 mM MgCl_2_ adjusted to pH of 7.4 with TEA-OH and 305 mOsm with TEA-Cl. The Cs-based internal solution contained 126 mM CsCl, 4 mM Mg-ATP, 1 mM EDTA, 10 mM EGTA, and 10 mM HEPES, adjusted to a pH of 7.4 with CsOH and an osmolarity of 290 mOsm with CsCl as needed. The K-based internal solution contained 126 mM KCl, 4 mM Mg-ATP, 1 mM EDTA, 10 mM EGTA, and 10 mM HEPES (pH adjusted with NaOH), adjusted to a pH of 7.4 with KOH and an osmolarity of 290 mOsm with KCl as needed.

We used a microperfusion system fed by gravity for rapid solution exchange during recording. This was comprised of 6-8 small diameter glass tubes aligned and adhered together and connected to inert tubing. Solution flow through each line was independently controlled via electronic valves. The glass tubes were immersed in the recording chamber using a micromanipulator. Once a stable recording was achieved, the cell is lifted from the bottom of the dish and carefully positioned at the outflow of the desired solution/pipe and solution exchange was achieved by manually moving the cell rapidly (within 1 sec) between pipes. There was a constant flow of the appropriate external solution applied via a larger outlet ensuring constant laminar flow. We applied 2 µM capsaicin in Na-based solution or 2 µM ω-Conotoxin GVIA (C-300, Alomone Labs) in a TEA-based solution to individual cells during recording.

Our previous analyses of DRG neurons in mice revealed the presence of three types of neurons based on size and the presence or absence of a prominent low threshold Ca_V_3 type current^18^. The smaller diameter cells (∼15 pF) included capsaicin responsive/TRPV1+ neurons. We biased recordings toward this cell population by primarily selecting smaller diameter neurons which were both positive and negative for tdTomato. We recorded neurons from *Cacna1b^fl/fl^*/*Trpv1^Cre/-^* (cKO), *Cacna1b^fl/fl^* (WT-fl) littermate controls and *Trpv1^Cre^*^/Cre^ (WT-cre) mice. Neurons were also grouped for analysis post-recording based on capsaicin responsiveness (Supplemental Fig. 2).

Whole cell voltage-gated calcium (Ca_V_) currents were recorded in TEA-based extracellular solution with 2 mM Ca^2+^ and elicited by depolarizing voltage steps form a holding potential of -80 mV. 2 µM ω-CgTx-GVIA was added to block Ca_V_2.2 currents. Leak currents were subtracted online using a P/-4 protocol. Currents were recorded using an Axopatch 200B amplifier, filtered at 5 kHz and digitized at 10 kHz (Axon Instruments 1440A). For 2 µM capsaicin responses, cells were moved to a Na-based extracellular solution, held at -60 mV, and exposed to capsaicin for 1 second. Currents were analyzed with Clampex 10.7 and custom lab software using Python 3.7 and pyABF^63^ and IPFX^64^ packages. Recording electrodes were fire-polished to between 2 and 5 MΩ, the average series resistance was 7.7 MΩ, and average cell capacitance 15.86 ± 0.54 pF. All recordings were obtained at room temperature.

### Behavioral assessments

All behavioral assessments were conducted by experimenters who were blind to the genotype of the mice. However, the hyperactivity phenotype of gKO mice and the edema and guarding associated with CFA injections were so consistent that the experimenter was able to distinguish these mice from WT or saline controls.

*Heat assessment*. Paw withdrawal responses were elicited by radiant heat applied using a Hargreaves equipment (Plantar Analgesia Meter IITC Life Science). Mice were placed in Plexiglas boxes on an elevated glass plate and allowed to habituate for at least 30 min prior to testing. A radiant heat source was positioned beneath the mouse paw and aimed using low-intensity visible light to the plantar surface. For all trials, laser idle intensity was 5% and active intensity was 50% of maximum. Trials were terminated if the mouse withdrew, shook, and/or licked their hindpaw following stimulation or if the cut off time, 30 s, was reached. The light source was manually turned off if any of these criteria were met. The response latency was measured to the nearest 0.01 s for each trial using the built-in timer corresponding to the duration of the high-intensity beam, see^4,5,10^.

*Mechanical assessment.* Mechanical responses were evoked and recorded with an automated von Frey plantar aesthesiometer (catalog #37550, Ugo Basile). Mice were placed in an elevated Plexiglas box with a wire mesh bottom and were allowed to habituate for at least 30 min prior to testing. The plantar surface of hind paws was assessed using a 15 s steady ramp of force ranging from 0 to 8 g for up to 90 s. The trial is automatically terminated when the paw is withdrawn or cutoff time is reached; force and reaction time are captured. Here, we report only the force associated with the withdrawal threshold^4,5,10^.

For both heat and mechanical assessment, three trials were conducted on each hindpaw for each mouse, with at least 1 min rest between trials. The average of 3 trials was used for analysis, and N values are reported as the number of mice. After baseline measures, mice were anesthetized with isoflurane for intradermal injection of inflammatory agents and returned to the testing chamber to recover. Males and females were tested separately in different sessions.

*Open Field*. Mice were tested a single time in an open field apparatus in dim light conditions. The apparatus consisted of an open field area of 20 x 20 x 14 in. Mice were recorded with an overhead video camera. Mice were tracked using Noldus EthoVision XT tracking software. After 30 min of habituation to the testing room, mice were placed in the center of the open field arena and allowed to freely explore for 5 min. Distance traveled was scored as a readout of activity. Males and females were tested separately in different sessions.

*Cold plate*. Sensitivity to cold was assessed using the Hot/Cold Plate Analgesia Meter from IITC, Inc. After 30 min of habituation to the testing room, mice were placed on a cold surface kept at - 5°C. Mice were allowed to freely move on the cold plate. A 30 s cut off was used for this experiment. Two cameras were used to record the behavior of the animals on the cold plate. Latency to response with a paw flinching, licking, shaking, or jumping was scored post hoc using the recorded videos. At least two independent experimenters blind to the genotype scored the videos with an interrater variability <5%. Males and females were tested separately in different sessions.

### Intradermal administration of inflammatory agents

For intradermal administration, mice were lightly anesthetized with isoflurane and a 30 G insulin syringe was used to inject 20 µL 0.1% w/v capsaicin in saline, 20 µL saline or 20 µL Complete Freund’s Adjuvant (CFA) into the plantar surface of the right hindpaw^4,5,10^.

### Statistical analysis

Data were assessed for normality using the Shapiro-Wilk test and for homogeneity of variance using Levene’s test. Based on these assumptions, appropriate parametric or non-parametric statistical tests were selected. For comparisons between two groups, Student’s t-test was used when data met assumptions of normality and equal variance. If variance was unequal, Welch’s t-test was performed. When data were non-normally distributed, the Mann-Whitney U test was used. For within-subject or paired comparisons, paired t-tests were used for normally distributed data, and the Wilcoxon signed-rank test was used when normality was violated. For multiple group comparisons, one-way or two-way ANOVA was used for normally distributed data, followed by Tukey’s HSD post hoc test when significant main effects or interactions were observed. If data violated normality or equal variance assumptions, the Kruskal-Wallis test with Dunn’s post hoc test was applied. For repeated measures designs, repeated measures ANOVA was used with Tukey’s HSD correction for post hoc pairwise comparisons. When random effects (e.g., repeated measures within subjects) needed to be accounted for, linear mixed-effects models were applied. All statistical analyses were performed using Python’s statsmodels and scipy packages. Data are presented as mean ± SEM, and we considered a *p* value of 0.05 as a measure of significance, but in all analyses, we show the actual *p* values together with information about the specific tests used.

## REFERENCES

1. Dolphin, A. C. & Lee, A. Presynaptic calcium channels: specialized control of synaptic neurotransmitter release. Nat. Rev. Neurosci. 21, 213–229 (2020).

2. Catterall, W. A. Voltage-Gated Calcium Channels. Cold Spring Harb. Perspect. Biol. 3, a003947 (2011).

3. Raingo, J., Castiglioni, A. J. & Lipscombe, D. Alternative splicing controls G protein– dependent inhibition of N-type calcium channels in nociceptors. Nat. Neurosci. 10, 285–292 (2007).

4. Salib, A.-M. N. et al. Interleukin-1α links peripheral CaV2.2 channel activation to rapid adaptive increases in heat sensitivity in skin. Sci. Rep. 14, 9051 (2024).

5. Salib, A.-M. N., Crane, M. J., Jamieson, A. M. & Lipscombe, D. Peripheral CaV2.2 Channels in the Skin Regulate Prolonged Heat Hypersensitivity during Neuroinflammation. eNeuro 11, (2024).

6. White, D. M. & Cousins, M. J. Effect of subcutaneous administration of calcium channel blockers on nerve injury-induced hyperalgesia. Brain Res. 801, 50–58 (1998).

7. Nebe, J., Vanegas, H. & Schaible, H.-G. Spinal application of ω-conotoxin GVIA, an N-type calcium channel antagonist, attenuates enhancement of dorsal spinal neuronal responses caused by intra-articular injection of mustard oil in the rat. Exp. Brain Res. 120, 61–69 (1998).

8. Lipscombe, D. & Andrade, A. Calcium Channel CaVα1 Splice Isoforms - Tissue Specificity and Drug Action. Curr. Mol. Pharmacol. 8, 22–31.

9. Zoidis, G., Sandoval, A., Pineda-Farias, J. B., Granados-Soto, V. & Felix, R. Anti-allodynic effect of 2-(aminomethyl)adamantane-1-carboxylic acid in a rat model of neuropathic pain: A mechanism dependent on CaV2.2 channel inhibition. Bioorg. Med. Chem. 22, 1797–1803 (2014).

10. DuBreuil, D. M. et al. Heat But Not Mechanical Hypersensitivity Depends on Voltage-Gated Ca _V_ 2.2 Calcium Channel Activity in Peripheral Axon Terminals Innervating Skin. J. Neurosci. 41, 7546–7560 (2021).

11. Beuckmann, C. T., Sinton, C. M., Miyamoto, N., Ino, M. & Yanagisawa, M. N-Type Calcium Channel α1B Subunit (CaV2.2) Knock-Out Mice Display Hyperactivity and Vigilance State Differences. J. Neurosci. 23, 6793–6797 (2003).

12. Gorman, K. M. et al. Bi-allelic Loss-of-Function *CACNA1B* Mutations in Progressive Epilepsy-Dyskinesia. Am. J. Hum. Genet. 104, 948–956 (2019).

13. Saegusa, H. et al. Suppression of inflammatory and neuropathic pain symptoms in mice lacking the N-type Ca2+ channel. EMBO J. 20, 2349–2356 (2001).

14. Nakagawasai, O. et al. Behavioral and neurochemical characterization of mice deficient in the N-type Ca2+ channel α1B subunit. Behav. Brain Res. 208, 224–230 (2010).

15. Caterina, M. J. et al. The capsaicin receptor: a heat-activated ion channel in the pain pathway. Nature 389, 816–824 (1997).

16. Caterina, M. J. et al. Impaired nociception and pain sensation in mice lacking the capsaicin receptor. Science 288, 306–313 (2000).

17. Cavanaugh, D. J. et al. Trpv1 reporter mice reveal highly restricted brain distribution and functional expression in arteriolar smooth muscle cells. J. Neurosci. Off. J. Soc. Neurosci. 31, 5067–5077 (2011).

18. Andrade, A., Denome, S., Jiang, Y.-Q., Marangoudakis, S. & Lipscombe, D. Opioid inhibition of N-type Ca2+ channels and spinal analgesia couple to alternative splicing. Nat. Neurosci. 13, 1249–1256 (2010).

19. Marangoudakis, S. et al. Differential Ubiquitination and Proteasome Regulation of CaV2.2 N-Type Channel Splice Isoforms. J. Neurosci. 32, 10365–10369 (2012).

20. Schiff, M. L. et al. Tyrosine-kinase-dependent recruitment of RGS12 to the N-type calcium channel. Nature 408, 723–727 (2000).

21. ISH Data :: Allen Brain Atlas: Mouse Brain. https://mouse.brain-map.org/.

22. Brain RNA-Seq – Brain RNA-Seq. https://brainrnaseq.org/.

23. Zheng, Y. et al. Deep Sequencing of Somatosensory Neurons Reveals Molecular Determinants of Intrinsic Physiological Properties. Neuron 103, 598–616.e7 (2019).

24. Harmonized cross-species cell atlases of trigeminal and dorsal root ganglia | Science Advances. https://www.science.org/doi/10.1126/sciadv.adj9173.

25. Gong, B., Murray, K. D. & Trimmer, J. S. Developing high-quality mouse monoclonal antibodies for neuroscience research – approaches, perspectives and opportunities. New Biotechnol. 33, 551–564 (2016).

26. Bernal Sierra, Y. A., Haseleu, J., Kozlenkov, A., Bégay, V. & Lewin, G. R. Genetic Tracing of Cav3.2 T-Type Calcium Channel Expression in the Peripheral Nervous System. Front. Mol. Neurosci. 10, (2017).

27. François, A. et al. The Low-Threshold Calcium Channel Cav3.2 Determines Low-Threshold Mechanoreceptor Function. Cell Rep. 10, 370–382 (2015).

28. Bourinet, E. et al. Silencing of the Cav3.2 T-type calcium channel gene in sensory neurons demonstrates its major role in nociception. EMBO J. 24, 315–324 (2005).

29. Lawson, J. J., McIlwrath, S. L., Woodbury, C. J., Davis, B. M. & Koerber, H. R. TRPV1 Unlike TRPV2 is Restricted to a Subset of Mechanically Insensitive Cutaneous Nociceptors Responding to Heat. J. Pain Off. J. Am. Pain Soc. 9, 298–308 (2008).

30. An experimental investigation of rundown … | Wellcome Open Research. https://wellcomeopenresearch.org/articles/9-250.

31. Hao, L.-Y., Kameyama, A., Kuroki, S., Nishimura, S. & Kameyama, M. Run-Down of L-Type Ca2+Channels Occurs on the α1Subunit. Biochem. Biophys. Res. Commun. 247, 844–850 (1998).

32. Wilhelm, J. C., Rich, M. M. & Wenner, P. Compensatory changes in cellular excitability, not synaptic scaling, contribute to homeostatic recovery of embryonic network activity. Proc. Natl. Acad. Sci. 106, 6760–6765 (2009).

33. O’Leary, T., Williams, A. H., Caplan, J. S. & Marder, E. Correlations in ion channel expression emerge from homeostatic tuning rules. Proc. Natl. Acad. Sci. 110, E2645–E2654 (2013).

34. Dhaka, A., Earley, T. J., Watson, J. & Patapoutian, A. Visualizing Cold Spots: TRPM8- Expressing Sensory Neurons and Their Projections. J. Neurosci. 28, 566–575 (2008).

35. Adams, D. J. & Berecki, G. Mechanisms of conotoxin inhibition of N-type (Cav2.2) calcium channels. Biochim. Biophys. Acta BBA - Biomembr. 1828, 1619–1628 (2013).

36. Pitake, S., Middleton, L. J., Abdus-Saboor, I. & Mishra, S. K. Inflammation Induced Sensory Nerve Growth and Pain Hypersensitivity Requires the N-Type Calcium Channel Cav2.2. Front. Neurosci. 13, 1009 (2019).

37. Klotz, U. Ziconotide--a novel neuron-specific calcium channel blocker for the intrathecal treatment of severe chronic pain--a short review. Int. J. Clin. Pharmacol. Ther. 44, 478–483 (2006).

38. Wallace, M. S. Ziconotide: a new nonopioid intrathecal analgesic for the treatment of chronic pain. Expert Rev. Neurother. 6, 1423–1428 (2006).

39. Murakami, M. et al. Modified sympathetic regulation in N-type calcium channel null-mouse. Biochem. Biophys. Res. Commun. 354, 1016–1020 (2007).

40. Yang, J. et al. Upregulation of N-type calcium channels in the soma of uninjured dorsal root ganglion neurons contributes to neuropathic pain by increasing neuronal excitability following peripheral nerve injury. Brain. Behav. Immun. 71, 52–65 (2018).

41. Chai, Z. et al. CaV2.2 Gates Calcium-Independent but Voltage-Dependent Secretion in Mammalian Sensory Neurons. Neuron 96, 1317–1326.e4 (2017).

42. Li, Y.-L. Stellate Ganglia and Cardiac Sympathetic Overactivation in Heart Failure. Int. J. Mol. Sci. 23, 13311 (2022).

43. Renganathan, M., Cummins, T. R. & Waxman, S. G. Contribution of Nav1.8 Sodium Channels to Action Potential Electrogenesis in DRG Neurons. J. Neurophysiol. 86, 629–640 (2001).

44. Blair, N. T. & Bean, B. P. Roles of Tetrodotoxin (TTX)-Sensitive Na+ Current, TTX-Resistant Na+ Current, and Ca2+ Current in the Action Potentials of Nociceptive Sensory Neurons. J. Neurosci. 22, 10277–10290 (2002).

45. Yoshida, S., Matsuda, Y. & Samejima, A. Tetrodotoxin-resistant sodium and calcium components of action potentials in dorsal root ganglion cells of the adult mouse. J. Neurophysiol. 41, 1096–1106 (1978).

46. Turrigiano, G., Abbott, L. F. & Marder, E. Activity-Dependent Changes in the Intrinsic Properties of Cultured Neurons. Science 264, 974–977 (1994).

47. Pratt, K. G. & Aizenman, C. D. Homeostatic regulation of intrinsic excitability and synaptic transmission in a developing visual circuit. J. Neurosci. Off. J. Soc. Neurosci. 27, 8268– 8277 (2007).

48. Black, J. A. et al. Abnormal expression of SNS/PN3 sodium channel in cerebellar Purkinje cells following loss of myelin in the taiep rat. NeuroReport 10, 913 (1999).

49. Waxman, S. G., Dib-Hajj, S., Cummins, T. R. & Black, J. A. Sodium channels and their genes: dynamic expression in the normal nervous system, dysregulation in disease states1. Brain Res. 886, 5–14 (2000).

50. Sensory neuron-specific sodium channel SNS is abnormally expressed in the brains of mice with experimental allergic encephalomyelitis and humans with multiple sclerosis | PNAS. https://www.pnas.org/doi/abs/10.1073/pnas.97.21.11598.

51. MacDonald, D. I. et al. A central mechanism of analgesia in mice and humans lacking the sodium channel NaV1.7. Neuron 109, 1497–1512.e6 (2021).

52. Jones, J. et al. Selective Inhibition of NaV1.8 with VX-548 for Acute Pain. N. Engl. J. Med. 389, 393–405 (2023).

53. Gingras, J. et al. Global Nav1.7 knockout mice recapitulate the phenotype of human congenital indifference to pain. PloS One 9, e105895 (2014).

54. Zakrzewska, J. M. et al. Safety and efficacy of a Nav1.7 selective sodium channel blocker in patients with trigeminal neuralgia: a double-blind, placebo-controlled, randomised withdrawal phase 2a trial. Lancet Neurol. 16, 291–300 (2017).

55. Hameed, S. Nav1.7 and Nav1.8: Role in the pathophysiology of pain. Mol. Pain 15, 1744806919858801 (2019).

56. Omari, S. A., Adams, M. J. & Geraghty, D. P. TRPV1 Channels in Immune Cells and Hematological Malignancies. Adv. Pharmacol. San Diego Calif 79, 173–198 (2017).

57. Mariotton, J., et al. TRPV1 activation in human Langerhans cells and T cells inhibits mucosal HIV-1 infection via CGRP-dependent and independent mechanisms. Proc. Natl. Acad. Sci. 120, e2302509120 (2023).

58. Bodó, E. et al. A Hot New Twist to Hair Biology: Involvement of Vanilloid Receptor-1 (VR1/TRPV1) Signaling in Human Hair Growth Control. Am. J. Pathol. 166, 985–998 (2005).

59. Bertin, S. et al. The ion channel TRPV1 regulates the activation and proinflammatory properties of CD4+ T cells. Nat. Immunol. 15, 1055–1063 (2014).

60. Cavanaugh, D. J. et al. Restriction of Transient Receptor Potential Vanilloid-1 to the Peptidergic Subset of Primary Afferent Neurons Follows Its Developmental Downregulation in Nonpeptidergic Neurons. J. Neurosci. 31, 10119–10127 (2011).

61. Korobkin, A. A. et al. Developmental Changes in the Expression of TRPV1 Channels in Autonomic Nervous System Neurons. Neurosci. Behav. Physiol. 43, 743–747 (2013).

62. Li, D., Li, Y., Tian, Y., Xu, Z. & Guo, Y. Direct Intrathecal Injection of Recombinant Adeno-associated Viruses in Adult Mice. JoVE J. Vis. Exp. e58565 (2019) doi:10.3791/58565.

63. Pyabf: Python Library for Reading Files in Axon Binary Format (ABF).

64. AllenInstitute/Ipfx. (Allen Institute, 2024).

