## Supplemental Figure 1 for "*Trpv1*-dependent *Cacna1b* gene inactivation reveals cell-specific functions of Ca_V_2.2 channels *in vivo*"

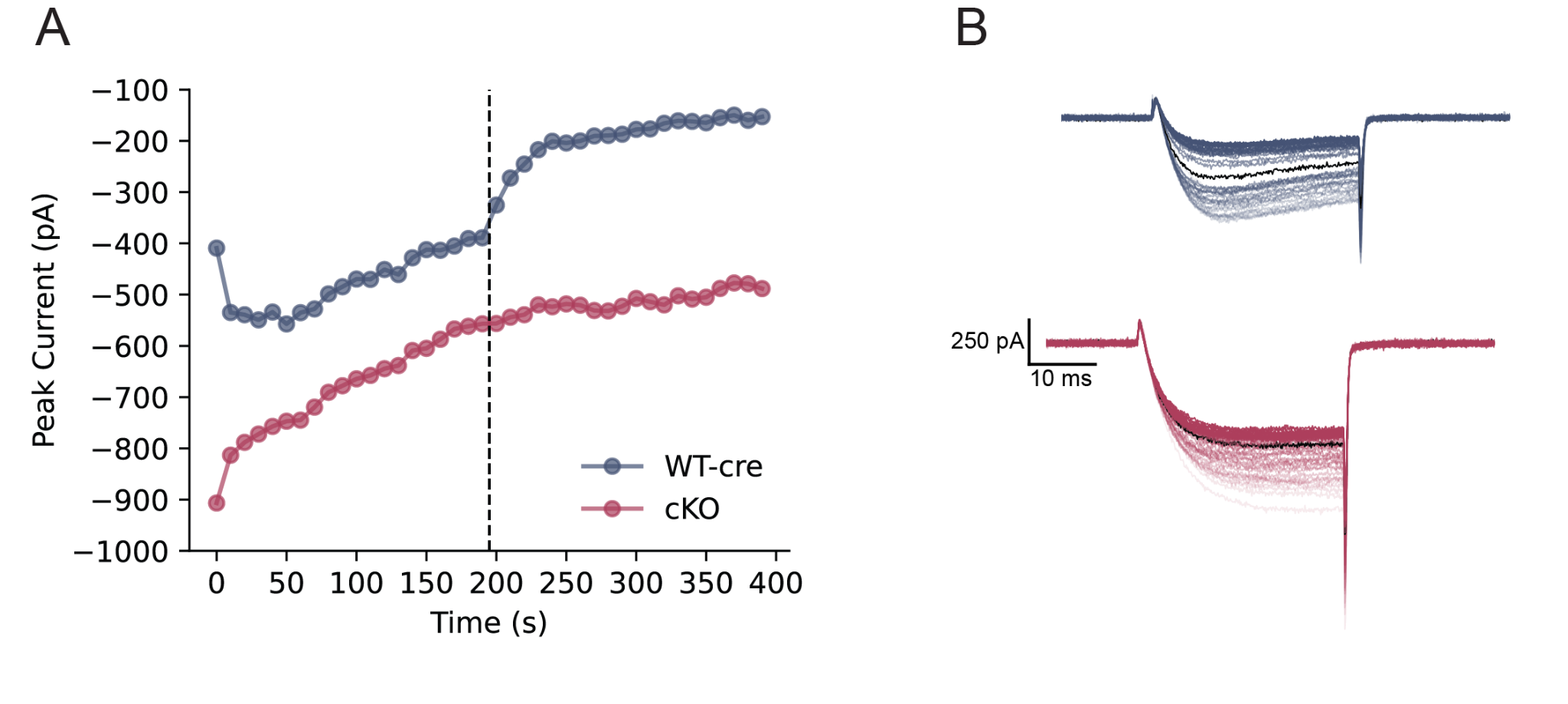

**Supplemental Figure 1. Calcium currents in *Trpv1*-lineage neurons from WT-fl and cKO mice exhibit rundown.** Whole cell Ca_V_ currents were recorded in *Trpv1*-positive (tdT+) DRG neurons isolated from *Trpv1^Cre/Cre^* (WT-cre, blue) and *Cacna1b^fl/fl^/Trpv1^Cre/-^* (cKO, red) mice. Ca_V_ currents were isolated using a TEA-based extracellular solution with 2 mM calcium as the charge carrier and elicited every 10 s by a depolarizing step to -10 mV from a holding potential of -80 mV. ***A***, shows the time course of peak current amplitude at -10 mV for a WT-cre and cKO recording before and after exposure to 2 µM ω-CgTx-GVIA (vertical dashed line. ***B***, the series of whole cell calcium current recordings from 0 to 400 sec (lighter to darker colors represent earlier and later times). The black trace in each series indicates the time of ω-CgTx-GVIA application. Ca_V_ currents from both WT-cre and cKO neurons exhibited a similar rate of rundown, but ω-CgTx-GVIA only inhibited WT-cre Ca_V_.
