## Supplemental Figure 2 for "*Trpv1*-dependent *Cacna1b* gene inactivation reveals cell-specific functions of Ca_V_2.2 channels *in vivo*"

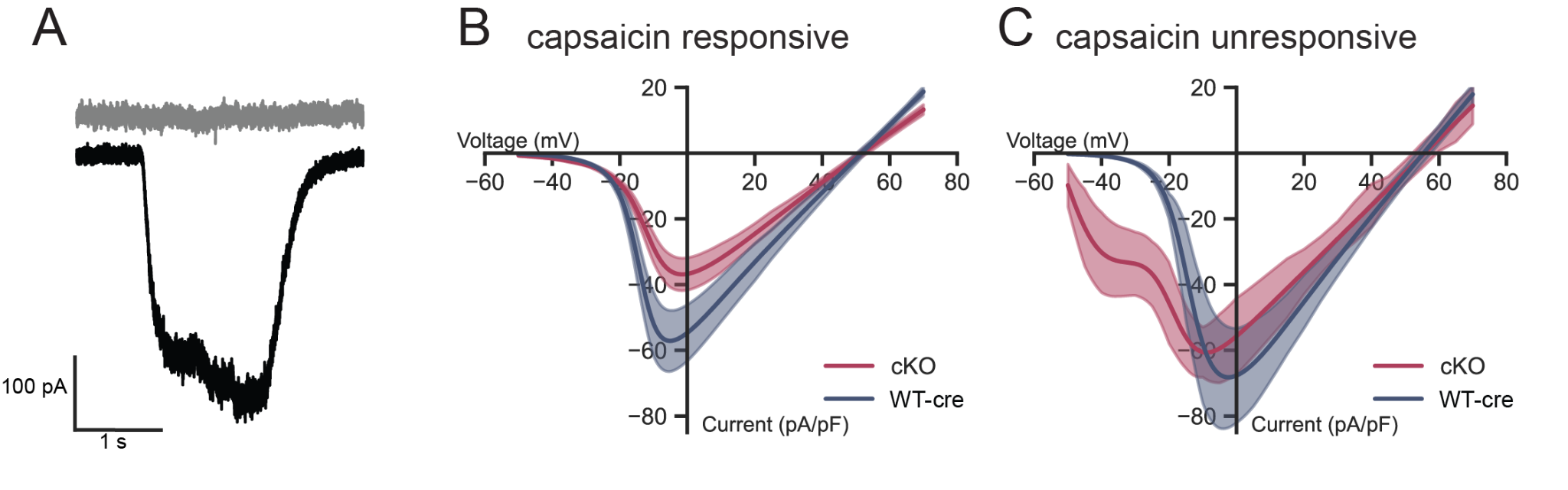

**Supplemental Figure 2. Whole cell calcium currents recorded from capsaicin-responsive and -nonresponsive cells.** Whole cell Ca_V_ currents were recorded in *Trpv1*-positive (tdT+) DRG neurons isolated from *Trpv1^Cre/Cre^* (WT-cre, blue) and *Cacna1b^fl/fl^/Trpv1^Cre/-^* (cKO, red) mice. Ca_V_ currents were isolated using a TEA-based extracellular solution with 2 mM calcium as the charge carrier and at the end of the experiment switched to a NaCl-based extracellular solution to test capsaicin-responsiveness using a 1 s exposure to 2 µM capsaicin. ***A***, Example recordings from capsaicin non-responsive (gray) and capsaicin-responsive (black) neurons. ***B***, Average peak Ca_V_ current densities for tdT+, capsaicin-responsive neurons from WT-cre (*n* = 13, blue) and cKO (*n* = 19, red) mice. Peak currents at 0 mV were significantly different between WT-cre and cKO (WT-cre | cKO: *t* = 2.227, *p* = 0.0336; Student’s t-test). ***C***, Average peak Ca_V_ current densities for tdT−, capsaicin nonresponsive cells from WT-cre (*n* = 5, blue) and cKO (*n* = 4, red). Peak currents at 0 mV were not significantly different between WT-cre and cKO (WT-cre | cKO: *t* = 0.4201, *p* = 0.6870; Student’s t-test).
